# Growth-driven displacement of protein aggregates along the cell length ensures partitioning to both daughter cells in *Caulobacter crescentus*

**DOI:** 10.1101/436394

**Authors:** Frederic D. Schramm, Kristen Schroeder, Jonatan Alvelid, Ilaria Testa, Kristina Jonas

## Abstract

All living cells must deal with protein aggregation, which can occur as a result of experiencing stress. In the bacteria *Escherichia coli* and *Mycobacterium smegmatis*, aggregates collect at the cell poles and are retained over consecutive cell divisions only in the daughter cell that inherits the old pole, resulting in aggregation-free progeny within a few generations. Here we have studied the *in vivo* kinetics of aggregate formation and clearance following heat and antibiotic stress in *Caulobacter crescentus*, which divides by a pre-programmed asymmetric cell cycle. Unexpectedly, we find that aggregates do not preferentially collect at the cell poles, but form as multiple distributed foci throughout the cell volume. Time-lapse microscopy revealed that under moderate stress, the majority of protein aggregates are short-lived and rapidly dissolved by the major chaperone DnaK and the disaggregase ClpB. Severe stress or genetic perturbation of the protein quality machinery results in long-lived protein aggregates, which individual cells can only clear by passing on to their progeny. Importantly, these persistent aggregates are neither collected at the old pole over multiple generations nor inherited exclusively by the old pole-inheriting stalked cell, but instead are partitioned between both daughter cells during successive division events in the same ratio. Our data indicate that this symmetric mode of aggregate inheritance is driven by the elongation and division of the growing mother cell. In conclusion, our study revealed a new pattern of aggregate inheritance in bacteria.

## Introduction

Exposure to various types of environmental stress results in the un- and misfolding of proteins, which poses a threat to continued survival. Sudden unfolding of native proteins, as well as interference of un- and misfolded proteins with partially folded species can create a functional deficit which impairs essential cellular processes and may lead to the death of the cell. All forms of life therefore rely on protein quality control systems to prevent accumulation of un- and misfolded proteins. The main participants in these systems are molecular chaperones and proteases that refold or degrade un- and misfolded proteins in order to maintain protein homeostasis as conditions fluctuate (Hartl et al., 2011). Acute stress can lead to exhaustion of the chaperoning and degradation capacity of the cell, resulting in aggregation of proteins that cannot be restored to their native state, either temporarily or indefinitely (Santra et al., 2018; Tyedmers et al., 2010). Sequestering un- or misfolded proteins into more inert particles has been proposed to lead to an immediate easing of the burden on the chaperone machinery, and can even be chaperone driven (Grousl et al., 2018; Ungelenk et al., 2016).

How cells cope with protein aggregates can differ depending on the type and amount of aggregated proteins present, but the highly conserved chaperone systems are responsible for effecting survival. Small heat shock proteins (sHSPs) associate with aggregated protein to maintain it in a refolding-competent state and can also promote the fusion of aggregates and facilitate their resolution (Coelho et al., 2014; Specht et al., 2011; Ungelenk et al., 2016). The cytoplasmic DnaK/Hsp70 chaperone and the ClpB/Hsp104 disaggregase bind insoluble aggregates and work in concert on their dissolution, returning proteins to their folded state (Glover and Lindquist, 1998; Goloubinoff et al., 1999). Additionally, cytosolic proteases contribute to aggregate resolution through the degradation of the constituent proteins (Heck et al., 2010; Tomoyasu et al., 2001).

If aggregates persist for a prolonged time and remain even after conditions improve, asymmetric distribution of insoluble deposits to daughter cells has been suggested to provide an effective means of sequestering un- and misfolded protein from a part of the population (Hill et al., 2016; Lindner et al., 2008; Vaubourgeix et al., 2015; Vedel et al., 2016; Winkler et al., 2010). For example, the budding yeast *Saccharomyces cerevisiae* retains aggregates in the mother cell both by active and passive mechanisms to generate aggregate-free daughter cells (Erjavec et al., 2007; Higuchi et al., 2013; Spokoini et al., 2012; Zhou et al., 2011). Although accumulation of protein aggregates has generally been associated with cell ageing and other pathology (Aguilaniu et al., 2003; Mogk et al., 2018; Nyström and Liu, 2014; Shcheprova et al., 2008), several recent studies suggest that the carriage of persistent protein aggregates may also confer a fitness advantage and promote survival (Govers et al., 2018; Saarikangas and Barral, 2015; Wallace et al., 2015).

So far, most studies addressing the *in vivo* dynamics of protein aggregate formation and clearance have been performed in eukaryotes. However, bacteria in particular frequently encounter stress conditions that perturb protein homeostasis, including heat, oxidative or antibiotic stress. In *Escherichia coli*, protein aggregates mostly accumulate at the chromosome-free polar regions of the cell (Kumar and Sourjik, 2012; Winkler et al., 2010). With successive cell divisions, this localization rapidly results in the production of aggregate-free cells (Govers et al., 2014; Lindner et al., 2008; Winkler et al., 2010). Similarly, slow-growing *Mycobacteria* were shown to collect irreversibly damaged proteins at the pole and distribute them asymmetrically to progeny, again resulting in aggregate-free daughter cells upon cell division (Fay and Glickman, 2014; Vaubourgeix et al., 2015). In both *E. coli* and *Mycobacterium*, the carriage of ancestral protein aggregates has been associated with a decline in growth rate (Lindner et al., 2008; Vaubourgeix et al., 2015; Winkler et al., 2010). However, a more recent study argues that *E. coli* cells inheriting protein aggregates along with components of the protein quality control machinery show an increased robustness to subsequent proteotoxic stress (Govers et al., 2018). Despite insight into the strategies and physiological consequences of aggregate distribution employed by *E. coli* and *Mycobacteria*, it remains poorly understood how other bacteria deal with protein aggregates in response to changing growth conditions. Of particular interest is how protein aggregation is handled by bacterial species possessing an intrinsically asymmetric life cycle, generating daughter cells with distinct cell fates.

The α-proteobacterium *Caulobacter crescentus* has long been a model organism of bacterial cell type differentiation as it undergoes asymmetric cell division. Each division cycle of *C. crescentus* yields two non-identical daughter cells, a motile, non-replicative swarmer cell, and a surface-attached and replication competent stalked cell (Curtis and Brun, 2010). As a free-living aquatic bacterium, it frequently encounters temperature fluctuations and other stresses that potentially threaten the folding state of the protein complement. A previous report has suggested that *C. crescentus* stalked cells undergo a slow replicative aging (Ackermann et al., 2003). While in yeast, replicative ageing has been attributed to an accumulation of protein aggregation in the mother cell (Aguilaniu et al., 2003; Coelho et al., 2014; Erjavec et al., 2007; Hill et al., 2014), the observed decline in the reproductive output of *C. crescentus* remains largely unexplained. Major components of the general chaperone machinery of *C. crescentus* have been described and their importance for stress resistance is known (Baldini et al., 1998; Da Silva et al., 2003; Schramm et al., 2017). However, to date it has not been studied how *C. crescentus* manages protein aggregation during its asymmetric life cycle, and the question persists if retention of protein damage in the stalked cell may explain the previously observed ageing effects.

In this study we have followed the dynamics of aggregate formation, clearance and inheritance following heat and antibiotic stress and recovery in *C. crescentus*. We demonstrate that protein aggregates form as multiple DnaK-attended foci throughout the cell volume and that the mechanism by which cells clear aggregates largely depends on the intensity of stress. Importantly, we show that in contrast to previously studied bacteria, persistent aggregates that form as a consequence of severe stress or genetic mutation do not sort to the old pole-containing stalked cell, but are instead distributed to both daughter cells at the same ratio over successive divisions.

## Results

### Heat and antibiotic stress induce relocalization of the *C. crescentus* chaperone machinery to foci of protein aggregation

In order to probe the dynamics and requirements of protein aggregation and resolution in *C. crescentus*, we constructed strains bearing fluorescently-tagged versions of the major heat shock chaperone DnaK and the bacterial disaggregase ClpB, at their respective native loci (Fig. 1A). Tagging these proteins did not result in viability defects at the optimal growth temperature of 30°C, although expressing the tagged version of ClpB correlated with a reduction in heat tolerance (Supporting Information Fig. 1). We found that DnaK tagged with the monomeric fluorescent protein mVenus (DnaK-mVenus) was diffusely localized throughout cells at a normal growth temperature of 30°C (Fig. 1B). To probe the localization of DnaK-mVenus at super-resolution, we imaged cells with stimulated emission depletion (STED) microscopy and found that the diffuse pattern of DnaK-mVenus at 30°C was representative of many small clusters of DnaK measuring 66 ± 23 nm (Fig. 1C). Upon exposure to a heat stress temperature of 40°C, DnaK-mVenus localization changed to a punctate pattern, suggesting that in *C. crescentus* protein aggregation is grouped into multiple foci that are distributed throughout the cell volume (Fig. 1B). Most cells contained between two and four DnaK-mVenus foci, while around 10% of the population harbored five or more foci (Fig. 1D). STED imaging of the foci formed during heat shock revealed large foci that measured 199 ± 56 nm (Fig. 1C). Determining the cellular position of DnaK-mVenus clusters showed that they occur with similar frequency along the cell length (Fig. 1E).

**Figure 1.**
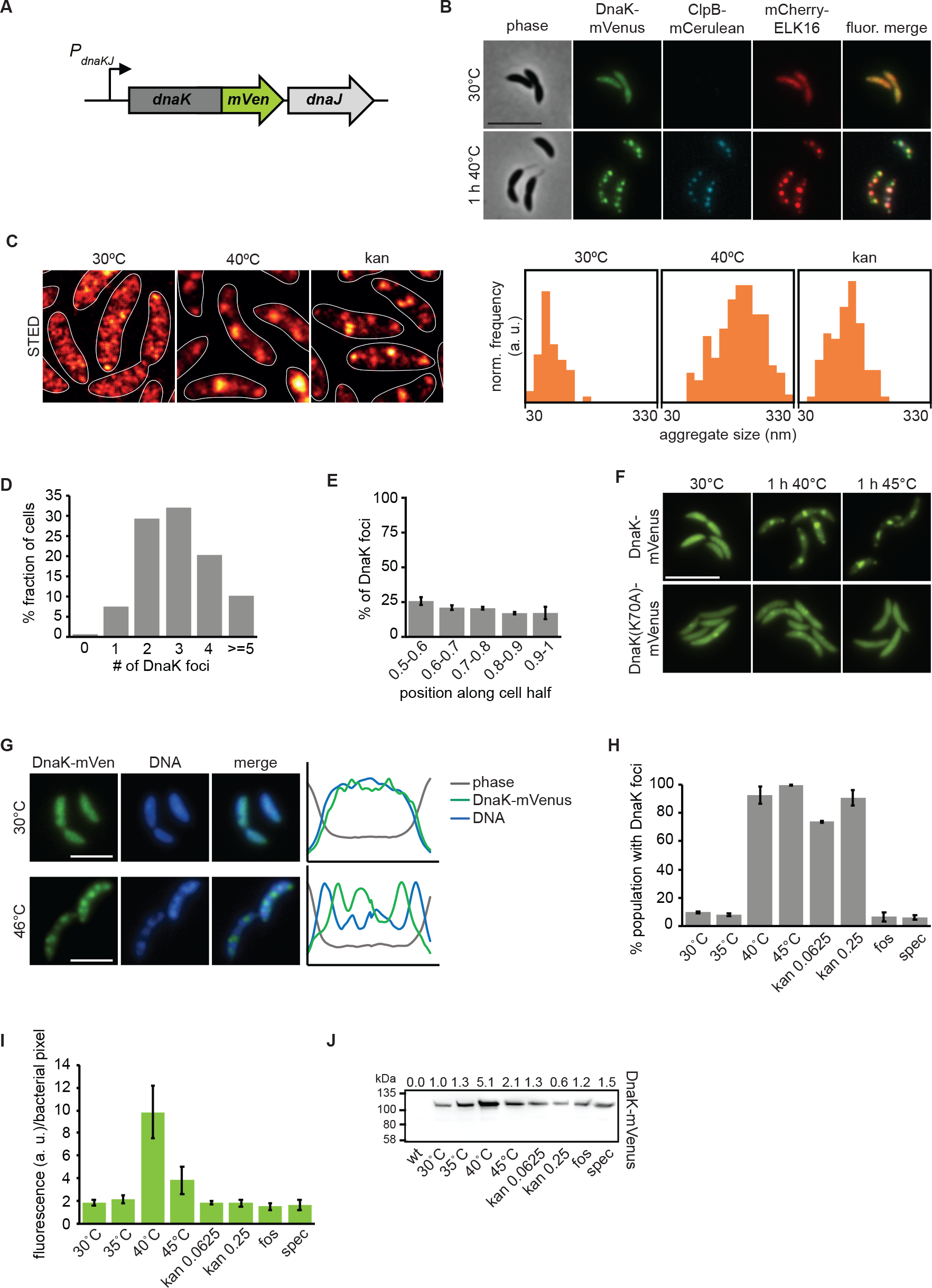
Stress induces relocalization of chaperone machinery to foci of protein aggregation. (A) Fluorescent fusions to proteins at their native locus are used to visualize the location of chaperones. (B) Microscopy of DnaK-mVenus, ClpB-mCerulean, and mCherry-ELK16 expressed at 30°C and after 1 h of heat shock at 40°C. (C) Representative images demonstrating DnaK-mVenus localization at super-resolution with STED imaging following incubation at 30°C, 1 h at 40°C, and after 1 h treatment with 0.25 μg/ml kanamycin. STED images are smoothed and cell outlines are shown in white. The histograms show the distribution of aggregate sizes, with all histograms showing the size range of 30-330 nm, and a bin width of 20 nm. Histograms show the size distribution of at least 60 aggregates. (D) Quantification of the number of aggregates per cell and (E) graph illustrating the position of the aggregates along five bins from midcell (0.5) to the cell pole (1) in cells exposed to 1 h of 40°C in liquid culture. Quantifications in (D) and (E) show the means of biological triplicates for which at least 196 cells and 685 aggregates each were analyzed, respectively. Error bars represent standard deviations. (F) Microscopy comparing localization of DnaK-mVenus with DnaK70A-mVenus at 30°C and after 1 h at 40°C. (G) Fluorescence microscopy demonstrating localization of DnaK-mVenus and the chromosome, stained with Hoechst 33258, after incubation at 30°C and 46°C. Line profiles (right) demonstrate fluorescence or phase contrast signal over the length of a representative cell from each treatment group. Signals are plotted as the percentage of the maximum value. Scale bar is 2.5 μm. (H) DnaK-mVenus foci formation in response to different stress conditions. Cultures of the DnaK-mVenus strain were exposed to the indicated temperature or treated with 0.0625 μg/ml or 0.25 μg/ml kanamycin, 20 μg/ml fosfomycin (fos), or 100 μg/ml spectinomycin (spec) for 1 h followed by visualization and quantification of the number of cells with DnaK-mVenus foci. Quantifications show the means of biological duplicates where 300 cells each were analyzed. Error bars represent standard deviation. (I) Quantification of DnaK-mVenus fluorescence intensity under different stress conditions. Cultures of the DnaK-mVenus strain were treated as in (H) followed by imaging and quantification of fluorescence intensity per bacterial pixel. Quantifications show the means of biological duplicates where at least 339 total cells each were analyzed from at least 10 independent images. (J) Quantification of protein levels of DnaK-mVenus under the conditions described in (H) and (I) by western blot. Band intensities are shown as average of two replicates above the western blot image.

Disabling ATP hydrolysis of DnaK by introducing a K70A mutation (Barthel et al., 2001) largely prevented DnaK from relocalizing to foci during heat stress, confirming that stress-induced DnaK relocalization is a function of its ATP-dependent foldase activity (Fig. 1F). To test whether the observed DnaK foci correspond to sites of protein aggregation, we used the aggregation-prone ELK16 peptide fused to mCherry as a marker for protein aggregation (Wu et al., 2011), and saw colocalization between DnaK-mVenus and mCherry-ELK16 foci after heat stress (Fig. 1B). Finally, co-localization experiments with the ClpB-mCerulean reporter showed that following exposure to heat stress ClpB attends mostly the same foci as DnaK-mVenus and mCherry-ELK16 (Fig. 1B), while no fluorescence was observed for the ClpB reporter at the normal growth temperature, in keeping with its heat shock-dependent expression (Simão et al., 2005). We note that ClpB foci were fewer and that some of the less intense foci of DnaK and mCherry-ELK16 were not attended by ClpB (Fig. 1B). This observation may be attributable to the function of ClpB in assisting refolding of large aggregates (Mogk et al., 2003b) or the reduced functionality of the ClpB-mCerulean fusion (Supporting Information Fig. 1). In addition to DnaK, ELK16 and ClpB, we have also tagged the small heat shock protein homolog CCNA_02341 (hereafter referred to as sHSP1). However, tagging this protein resulted in increased heat sensitivity, the formation of atypically large fluorescent clusters, and cell division defects during mild heat stress (Supporting Information Fig. 1). Based on these experiments we decided to utilize DnaK-mVenus for the visualization of aggregate localization throughout this study.

To determine the relationship between aggregate foci and the *Caulobacter* chromosome, we stained the DNA using Hoechst 33258 in the DnaK-mVenus reporter strain sampled at normal temperature as well as after heat shock (Fig. 1G). Surprisingly, aggregate formation during heat shock was accompanied by a change in the spatial organization of the DNA from an evenly dispersed to a patchy pattern. Moreover, the locations occupied by protein aggregates corresponded to regions of reduced DNA staining intensity. These data suggest that in contrast to *E. coli* and other bacteria, in which the chromosome pushes aggregates to the cell poles (Coquel et al., 2013; Lindner et al., 2008; Winkler et al., 2010), in *Caulobacter* protein aggregates appear to displace the chromosome.

Aggregate formation was observed at heat shock temperatures of 40°C and 45°C (Fig. 1H), but also upon exposure to sublethal concentrations of kanamycin, which is known to induce protein aggregation through mistranslation in *E. coli* (Kohanski et al., 2008). STED imaging revealed that kanamycin-induced aggregates were smaller than those induced by heat shock, forming as foci measuring 127 ± 39 nm (Fig. 1C). No aggregation foci were observed following treatment with the antibiotic fosfomycin, which blocks production of peptidoglycan precursors, nor for spectinomycin (Fig. 1H), a ribosome-binding aminoglycoside that does not cause mistranslation (Kohanski et al., 2008).

Finally, our reporter strains also allowed us to quantify the induction of *dnaK* expression following stress exposure by determining the fluorescence intensity per bacterial pixel in different conditions (Fig. 1I). Exposure to 40°C for one hour induced a four-fold increase in DnaK levels, while a temperature of 45°C induced a two-fold increase within one hour, which we confirmed by western blotting (Fig. 1J). Although strong aggregation was present, lower induction of DnaK at 45°C and following kanamycin treatment is possibly due to lower translation capacity in these conditions (Chen et al., 2017). Treatment with fosfomycin or spectinomycin did not increase DnaK levels, indicating that the inhibitory effects of these antibiotics act independently of protein aggregation and the heat shock response in *C. crescentus* (Fig. 1I, J). Altogether, these data establish that protein aggregation occurs at multiple sites in *C. crescentus* and that the major chaperones DnaK and ClpB are recruited to these sites.

### Proteins governing diverse processes comprise aggregate foci in *C. crescentus*

Having established a system for visualizing total cellular aggregation, we next asked what proteins might be un- or misfolding, thereby recruiting DnaK to foci of aggregation. In order to address this question, we isolated insoluble detergent-resistant proteins from wild type *C. crescentus* grown at 30°C or exposed to a stress temperature of 45°C for 1 hour, subjected them to mass spectrometry, and identified 133 proteins enriched specifically in the aggregated fraction during heat stress (Fig. 2A, Supporting Information Tab. 2). Comparing the abundance of aggregate-enriched proteins sorted by functional category (Fig. 2B), showed that proteins belonging to various cellular processes become associated with the aggregate foci, including 54 essential proteins (Supporting Information Tab. 2). The most abundant protein in the aggregated fraction was sHSP1, which shares 54% and 40% amino acid sequence homology with the *E. coli* small heat shock proteins IbpA and IbpB, respectively. This result is consistent with a previously described role of small heat shock proteins in binding and maintaining un- or misfolded proteins in a disaggregation-ready state (Lindner et al., 2008; Mogk et al., 2003a; Strozecka et al., 2012).

**Figure 2.**
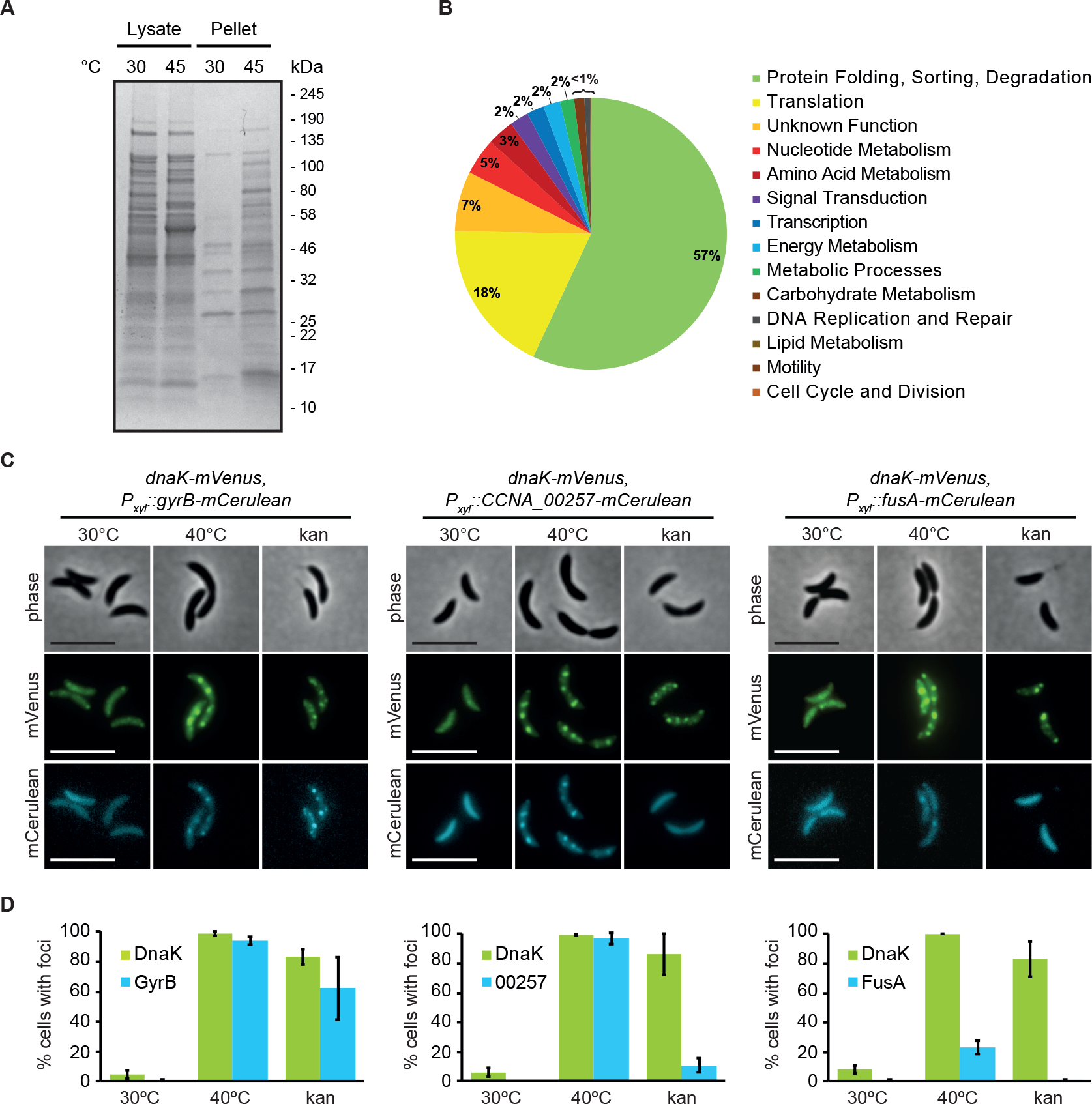
Proteins governing diverse processes aggregate in *C. crescentus*. (A) Coomassie stained protein gel of soluble and detergent-resistant aggregated fractions following heat stress. Wild type cultures at 30°C and 1 h at 45°C were collected and soluble and insoluble fractions were separated by lysis and centrifugation. (B) Abundance of aggregate-enriched proteins sorted by functional category. Functional categories were assigned to proteins identified in (A) according to KEGG gene ontology, and the enrichment of aggregation-prone proteins belonging to each category was determined to obtain relative abundance. (C) Localization pattern of the aggregation-prone proteins GyrB, CCNA_00257, and FusA at 30°C and 40°C. Expression of the fusion proteins was induced for 3 h at 30°C, followed by 1 h exposure to 40°C or 0.25 μM kanamycin. Scale bar is 5 μm. (D). Quantification of stress induced relocalization of endogenous aggregating proteins and DnaK-mVenus shown in (C). Quantifications show the means of biological duplicates where at least 152 cells each were analyzed. Error bars represent standard deviation.

To verify the identity of aggregated proteins detected by mass spectrometry, we selected the proteins GyrB, FusA, and the predicted homocysteinase CCNA_00257 to monitor their subcellular localization when exposed to heat and kanamycin stress (Fig. 2C). All three proteins were confirmed to condense into aggregate foci colocalizing with DnaK when exposed to stress (Fig. 2C). Interestingly, in contrast to GyrB, which relocalized in the majority of cells during all exposures, the number of cells with CCNA_00257 and FusA aggregates differed depending on the stress signal (Fig. 2D). While heat stress induced foci formation by CCNA_00257 in nearly all cells, kanamycin treatment resulted in foci formation in only 15% of the population, indicating that CCNA_00257 is less sensitive to kanamycin-induced effects than GyrB. FusA only relocalized into aggregate foci in 20% of the population during heat stress, again indicating that individual proteins are destabilized by different stress exposures. Collectively our data show that proteins belonging to diverse cellular processes become members of the aggregate foci observed in *C. crescentus* and that the composition of aggregates is dependent on the stress condition.

### Contribution of major chaperones and proteases to stress resistance and aggregate dissolution

In order to maintain cellular protein homeostasis, chaperones and proteases forming the protein quality control machinery must cooperate in protein folding, aggregate resolution, and degradation of un- and misfolded proteins. Through a series of deletion mutants we probed the contribution of ClpB, sHSP1, the other small heat shock protein homolog CCNA_03706 (hereafter referred to as sHSP2), and the protease Lon to aggregate resolution and stress resistance in *C. crescentus*.

In accordance with low expression during normal growth temperature, deletion of *clpB*, *shsp1*, and *shsp2* had no discernible effect at 30°C (Fig. 3A, B). Upon heat stress, the viability of cells lacking ClpB was drastically reduced, as has been demonstrated previously (Simão et al., 2005). The number and distribution of DnaK foci was similar to that observed in the presence of ClpB, however even when challenged by a sublethal heat shock cells were still unable to resolve aggregated protein deposits, which instead persisted over many generations even after release from stress (Supporting Information Fig. 2). In contrast to this pronounced phenotype, absence of *shsp1* or *shsp2* alone or in combination had no effect on resistance to heat treatment (Fig. 3A). The overall pattern of protein aggregation was unchanged in the absence of the sHSPs, forming still as distributed punctate foci, and aggregates were resolved within the same time frame as the parental strain (Fig. 3B). Basal levels of DnaK as well as levels induced by heat stress were similar in the absence of the sHSPs (Fig. 3C), indicating that additional DnaK is not compensating for the activity of these proteins and that although sHSP1 comprises nearly half of the aggregated protein fraction, both sHSPs are largely dispensable for tolerating acute heat stress under the stress conditions tested.

**Figure 3.**
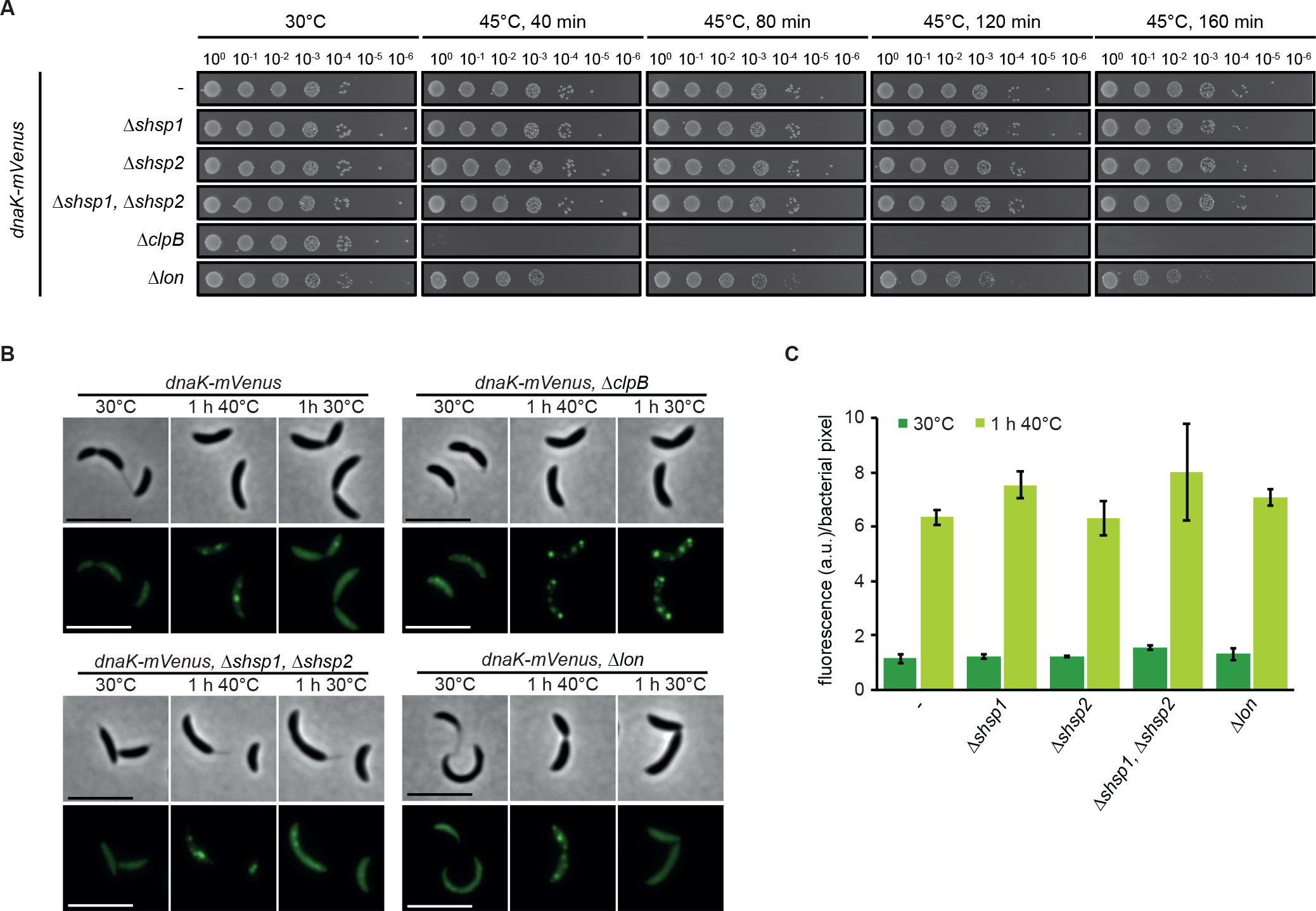
Contribution of chaperones and proteases to stress resistance. (A) Spot assays comparing colony formation of wild type, the *dnaK::dnaK-mVenus* strain and its derivative strains harboring chaperone and protease knockouts grown at 30°C without stress treatment or after 40, 80, 120 or 160 minutes of incubation at 45°C. (B) DnaK-mVenus localization in the same strains as in (A) grown at 30°C or after heat shock in liquid culture at 40°C for one hour (1 h 40°C panels). After 40°C heat shock, cells were permitted to recover at 30°C for one hour (1 h 30°C panels), during which aggregate dissolution was monitored by time lapse microscopy. (C) Absolute measurements of DnaK-mVenus fluorescence during growth at 30°C and after one hour heat shock at 40°C. Quantifications show the means of biological duplicates where at least 409 cells each were analyzed from 10 independent images.

Proteases support foldases in maintaining protein homeostasis by degrading un- or misfolded proteins, and combined loss of proteases has been shown to induce strong defects in protein quality control (Kanemori et al., 1997). Lon is generally viewed as the major protein quality control protease in *E. coli* (Rosen et al., 2002; Van Melderen and Aertsen, 2009), and has a role in cell cycle control and responding to unfolded protein in *C. crescentus* (Jonas et al., 2013). Cells lacking Lon exhibited only slight reduction in viability during heat stress (Fig. 3A), and both *dnaK-mVenus* induction and the pattern of aggregation occurred as when Lon was present. Additionally, in the absence of Lon cells were able to completely dissolve protein aggregates within a similar time frame as the parental strain (Fig. 3B, C). Thus, the Lon protease is not required for resolving protein aggregation under these stress conditions.

Together our data demonstrate that clustering of protein aggregation into several smaller foci and the recruitment of DnaK to these sites does not depend on other heat shock proteins. Furthermore, our data confirm in *C. crescentus* that resolution of protein aggregates requires the disaggregase ClpB and the chaperone DnaK, and that despite the heat shock induction of sHSPs and Lon, these factors are dispensable under the conditions tested.

### Aggregate clearance through dissolution or dilution is dictated by stress severity

As we have identified aggregate composition as well as chaperone contributions to aggregate dissolution, we next wanted to determine how *C. crescentus* responds to protein aggregation provoked by different stress intensities and how aggregates are cleared following shift to non-stress conditions. We therefore used fluorescence time-lapse microscopy to follow aggregate formation and resolution dynamics during different stress exposures.

To analyze the formation of aggregates following a temperature upshift, we transferred cultures grown at the normal temperature of 30°C to agarose pads and placed them into the imaging system pre-heated to either 30, 40, or 44°C, and monitored the localization pattern of DnaK-mVenus as a proxy for aggregate formation over time (Fig. 4A). Under both heat shock conditions, we observed rapid formation of an average of 3.7 aggregates per cell within 10 min of heat exposure (Fig. 4A, B, Supporting Information Movie 1). During incubation at 40°C, cells grew into microcolonies which increased in area at a rate similar to those in non-stress conditions, and aggregate formation and dissolution appeared highly dynamic (Fig. 4B, C). After their initial appearance the foci number per cell was quickly reduced until reaching a plateau of approximately one aggregate per cell by approximately two hours. We observed that the lifespan of most aggregates was less than 10 min (Fig. 4D). Only 23% of aggregates persisted for more than two hours (Fig. 4D), demonstrating that the aggregate numbers following two hours of exposure to 40°C represent a steady state of formation and disappearance of mostly short-lived aggregates as cells grow, with a minority of persistent aggregates emerging. We attribute the initial net reduction of aggregate number per cell (Fig. 4B) to continued growth and division and rapid induction of heat shock gene expression, which allows cells a greater capacity to cope with higher amounts of unfolded protein present at the new elevated temperature (Fig. 4E).

**Figure 4.**
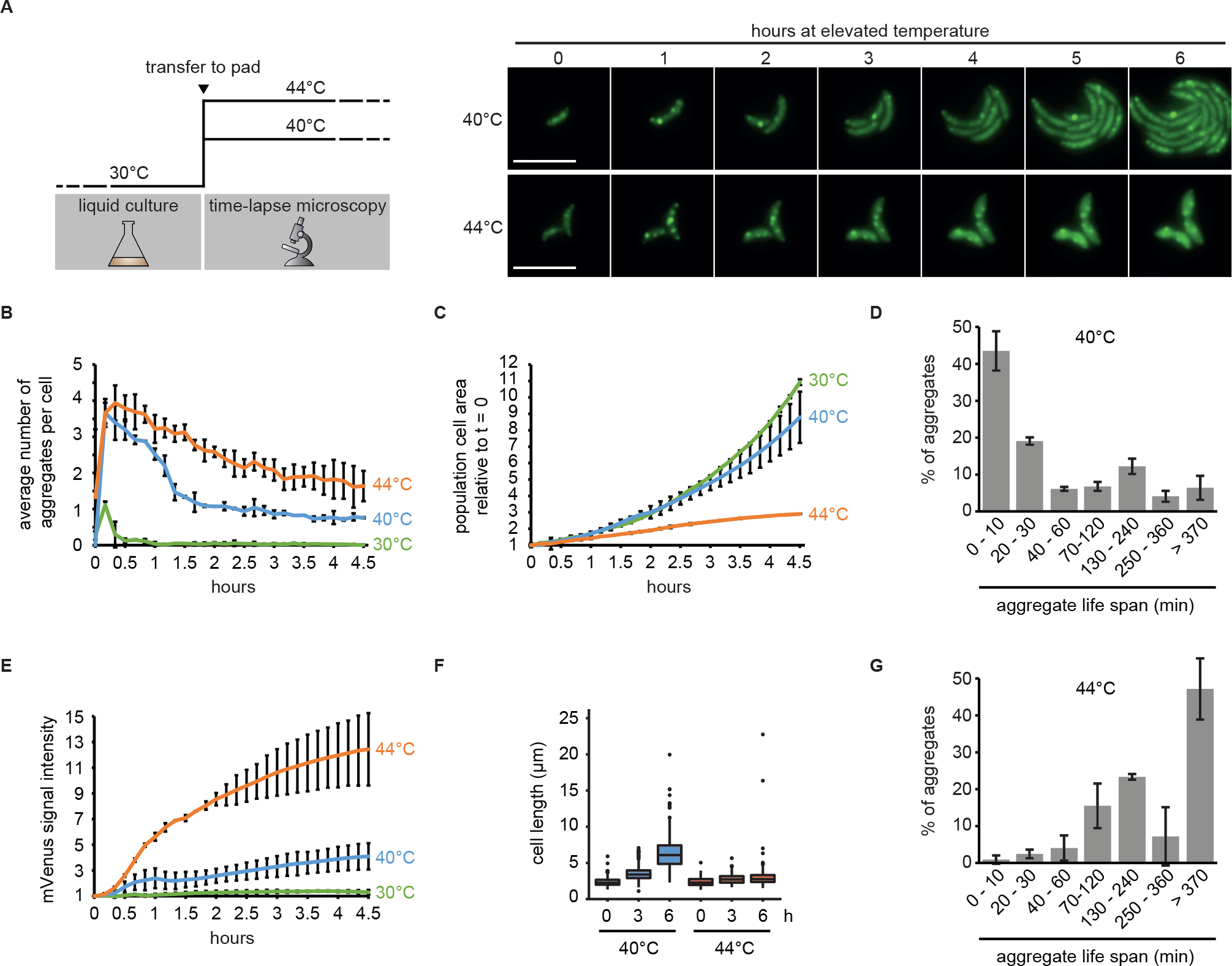
Growth and protein aggregation dynamics during sustained heat stress. (A) Schematic of upshift experiments (left), and representative images following upshift to 40°C or 44°C over a six hour period. (B) Quantifications of fluorescence time-lapse microscopy images showing the average number of aggregates per cell over time in cell populations continuously grown at 30, 40 and 44°C. Note that after transfer to 30°C a minor fraction of dim and short-lived DnaK-mVenus foci formed. (C) Total cell area increase during exposure to 30, 40, or 44°C relative to the first time point (t=0). Quantifications in (B, C) show the means of biological triplicates for which at least 8 microcolonies each were analyzed. Error bars represent standard deviations. (D) Quantification of aggregate life span in cells continuously exposed to 40°C. Aggregates present or emerging in the first 300 min of a fluorescence time-lapse movie were tracked until 400 min. Images were acquired every 10 min. Quantifications show the means of biological triplicates for which at least five microcolonies and 89 aggregates each were analyzed. Error bars represent standard deviations. (E) DnaK-mVenus fluorescence intensity per bacterial pixel over time in cells grown at 30, 40, or 44°C normalized to the first time point (t=0). The same cells as in (B, C) were quantified. (F) Quantification of population cell lengths after 0, 3, and 6 hours continuous exposure to 40, or 44°C. The cell lengths of two biological replicates were pooled and populations representing at least 504 measurements are shown. (G) Quantification of aggregate life span in cells continuously exposed to 44°C. Aggregates present at 0 min were tracked until 480 min (new aggregates did not arise). Images were acquired every 10 min. Quantifications show the means of biological duplicates for which at least 14 microcolonies and 63 aggregates each were analyzed. Error bars represent standard deviations.

Although microcolonies grew at a relatively normal rate, we observed that temperatures near 40°C resulted in abnormal filamentous growth phenotypes, indicative of cell division defects (Fig. 4F), consistent with an earlier study (Heinrich et al., 2016). At a temperature of 44°C cell growth was essentially arrested (Fig. 4A, C) while DnaK-mVenus levels drastically increased over time, indicating strong and persistent induction of heat shock gene expression (Fig. 4E). We observed that DnaK foci slowly dissolved until only 40% remained after 6 to 8 hours of exposure to 44°C, and only rarely observed the formation of new aggregates at this temperature (Fig. 4B, G). This observation indicates that a fraction of aggregates can be resolved by the highly abundant chaperones, or that DnaK is released from persistent aggregates under this severe continuous stress.

To monitor the resolution of aggregates as cells recovered from heat stress, we exposed cells growing in liquid culture to 40, 42, 44 or 46°C for one hour, followed by transfer to agarose pads for imaging at 30°C (Fig. 5A, Supporting Information Fig. 3). During recovery from these temperatures some or all of the stress-treated population remained capable of resuming growth (Fig. 5B) and reducing the average number of aggregates per cell (Fig. 5C). How quickly the average number of aggregates was reduced and whether this was accomplished by aggregate dissolution or dilution within a growing microcolony was dependent on the severity of the heat stress. Growth resumption and aggregate reduction were equally fast in cells exposed to 40 and 42°C (Fig. 5B, C), where all protein aggregates were completely dissolved within one hour (Fig. 5C), leading to aggregate-free microcolonies (Fig. 5A, D). In contrast, cells recuperating from exposure to 46°C showed a strong growth delay and decreased ability to clear aggregates, and were heterogenous both in aggregate resolution and the ability to produce normally growing progeny (Fig. 5A, B, D, E). Only 25% of cells were able to produce microcolonies of normally sized progeny (Fig. 5D), in which the average aggregate number per cell was notably reduced (Fig. 5C, E). The remaining 75% of the population were either unable to resume growth within 4.5 hours (32%) or resumed growth with varying degrees of division defects (43%) (Fig. 5D). Importantly, although most individuals within the forming microcolonies became eventually aggregation-free, a fraction of offspring maintained aggregates. Many of these aggregates originated from the heat exposed mother cell and persisted for the duration of imaging. However, we also observed that a fraction of new shorter-lived aggregates formed during recovery (Fig. 5F). Based on these analyses we conclude that a portion of aggregates formed during 46°C exposure persist over several generations and are cleared from individual cells mainly through dilution within the microcolony rather than dissolution. A similar behavior was also observed after exposure to 40°C for 10 min in the Δ*clpB* strain, where despite efficient clearance of aggregates from individual cells, all aggregates were long-lived (Supporting Information Fig. 2).

**Figure 5.**
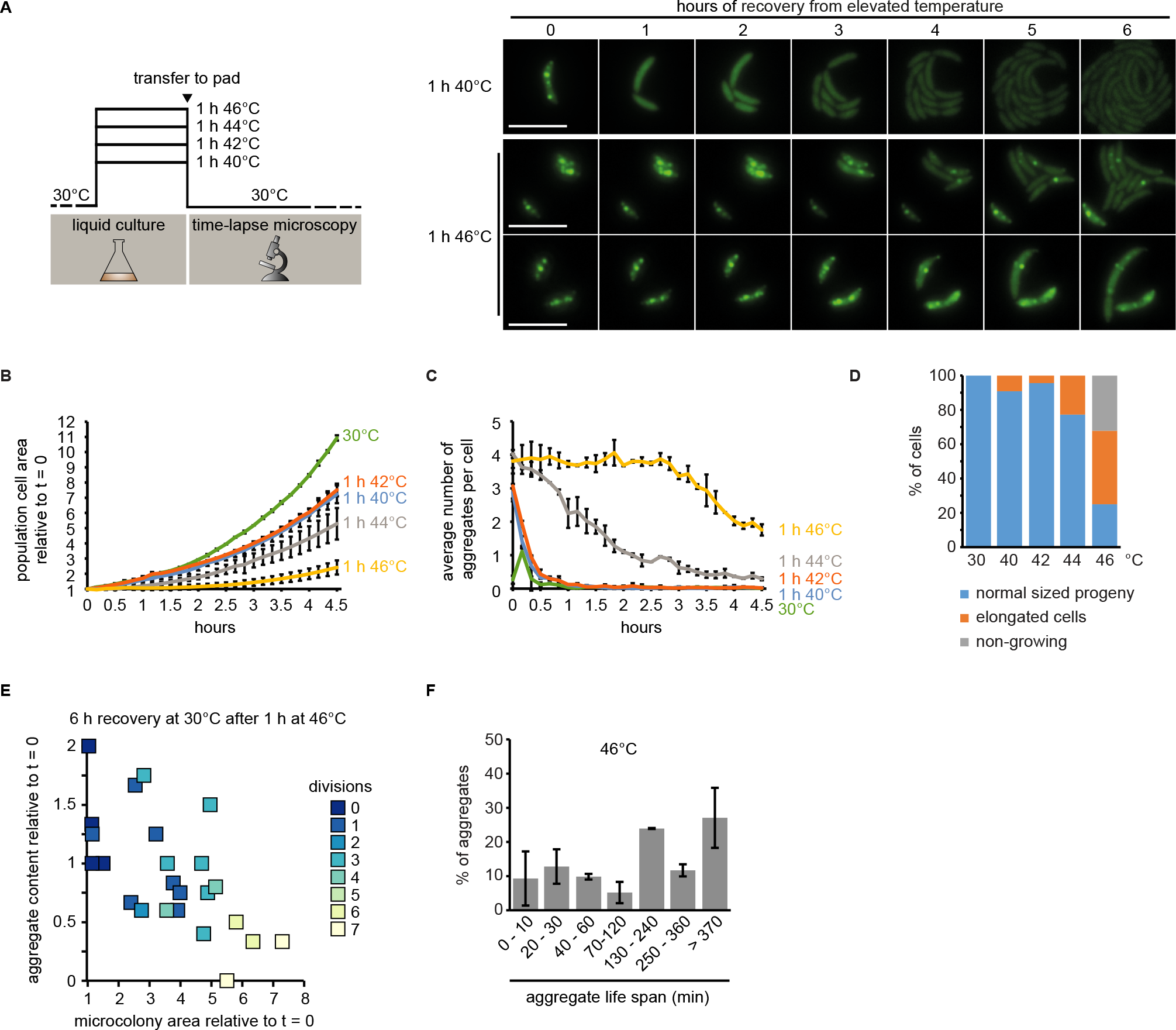
Growth and aggregate clearance following stress recovery. (A) Schematic of recovery experiments (left) and representative images from recovery after exposure to 40°C and 46°C for one hour. Two panels are shown for the 46°C condition to represent the diversity of cell fates after stress release. (B) Quantifications of fluorescence time-lapse microscopy showing the average number of aggregates per cell in cell populations recovering from one hour of exposure to 40, 42, 44 and 46°C. (C) Total cell area increase over time after release from one hour exposure to 40, 42, 44 and 46°C relative to the first time point (t=0). Quantifications in (B, C) show the means of biological duplicates for which at least 9 microcolonies each were analyzed. The data on cells continuously grown at 30°C are the same as in Fig. 4. Error bars represent standard deviations. (D) Quantification of cell fates during continuous growth at 30°C, or recovery at 30°C after 1 h of exposure to 40, 42, 44 and 46°C. Quantifications are based on tracking at least 21 cells/microcolonies per duplicate condition. Cell/microcolony properties were quantified when the average total area of the population increased to four times the initial area. Growing cells/microcolonies were defined as those that at least doubled in area. Elongated cells or microcolonies containing those were considered as such when the average cell length in the microcolony was at least 1.5 times higher than the average cell length of a population grown to four times the cell area under continuous 30°C conditions. (E) Scatter plot representing relative area increase, relative aggregate number normalized to the amount present at stress release and the number of divisions of 24 cells and their potential progeny after six hours (the time at which the average area of the population has quadrupled) at 30°C on pad after release from one hour of exposure to 46°C. (F) Quantification of aggregate life span in cells recovering from exposure to 1 h 46°C forming microcolonies of mostly normally sized cells (D). Aggregates present or emerging in the first 400 min of a fluorescence time-lapse movie were tracked until 500 min. Images were acquired every 10 min. Quantifications show the means of biological duplicates for which at least eight microcolonies and 54 aggregates each were analyzed. Error bars represent standard deviations.

When monitoring the recovery of cells following exposure to 44°C we again observed that aggregates are cleared through a combination of dissolution and dilution (Supporting Information Fig. 4). However, in contrast to the 46°C condition, the fraction of cells able to produce normally sized progeny was larger (77%) (Fig. 5F), cells resumed growth earlier (Fig. 5B) and nearly all cells were free of protein aggregates by 4.5 hours after exposure (Fig. 5C).

Taken together our results demonstrate that the severity of heat stress determines the way by which aggregates are cleared from the cell. Following exposure to moderate stress, virtually all aggregates are rapidly dissolved by the protein quality control machinery. By contrast, aggregate clearance following severe stress depends more on dilution during cell division. Consequently, an inability to resume growth and cell division following such severe stress prevents successful aggregate clearance.

### The aggregate load does not sort to swarmer or stalked daughter cells in *C. crescentus*

Asymmetric inheritance of protein aggregates has been proposed to underpin the senescence of aggregate retaining cells and the rejuvenation of aggregate free cells (Ackermann et al., 2003; Coelho et al., 2014; Lindner et al., 2008; Shapiro et al., 2002; Winkler et al., 2010). While asymmetric inheritance in *E. coli* and *Mycobacteria* takes place through collection of aggregates at the poles, in budding yeast asymmetric inheritance is achieved through retention in the much larger mother cell. Since *C. crescentus* has a pre-programmed asymmetric division cycle yielding a bigger stalked/old pole and a smaller swarmer/new pole cell, we sought to understand how persistent aggregates are inherited in this organism.

To study aggregate inheritance when only dilution can be used as a means of aggregate clearance, we made use of the Δ*clpB* background to investigate the distribution of aggregates following division. A short heat shock of 10 minutes at 40°C induced formation of persistent protein aggregates in this strain that were distributed throughout the cell volume, while the ability to grow and divide was preserved (Supporting Information Fig. 2), therefore we followed the inheritance of these aggregates over consecutive cell divisions by time lapse microscopy (Fig. 6A). A kymograph normalized to the summed length of all descendants from one individual cell shows that aggregates are relatively static and rarely change their cellular position between two cell division events (Fig. 6B). However, establishment of new cell boundaries as a consequence of cell division affected the relative cellular position of aggregates; for example, an aggregate located at midcell of a stalked cell becomes located closer to the new pole in the daughter cell following division (Fig. 6A, B). To analyze the positional change of aggregates more quantitatively, we determined the frequency by which aggregates obtain a new cellular position in the daughter stalked cell after cell division, as a function of their original cellular position (Fig. 6C, Supporting Information Fig. 5). Consistent with the kymograph analysis, these data show that aggregates located at midcell in the mother cell were likely to change relative position (71%) during cell division to become closer to the new pole of the stalked daughter cell. Aggregates between the old pole and midcell (classified as old pole half) either maintained their original relative position (64%) or also obtained a new relative position close to the new pole (34%). By contrast, aggregates located at the pole of the mother cell generally maintained their position during cell division (90%) and changed to a position closer to the new pole in only 10% of cases. Importantly, we only very rarely observed that aggregates located outside pole regions became situated closer to the old pole as a consequence of cell division (2% and 3% of those in the old pole half or at midcell, respectively), indicating that essentially all aggregates that obtain a new relative position in the daughter stalked cell after division, are situated closer to the new pole. We explain this passive “movement” of aggregates towards the new pole with the growth mode of *C. crescentus*, in which cells grow along the cell length, excluding the poles (Aaron et al., 2007; Lambert et al., 2018). Consequently, the fraction of aggregates originally formed at the poles will remain there over generations due to the absence of cell elongation in this area. Therefore, the majority of aggregates, while static, are displaced in proportion to the elongation of the cell and maintain their relative position until a division event sets new cellular boundaries (Fig. 6D).

**Figure 6.**
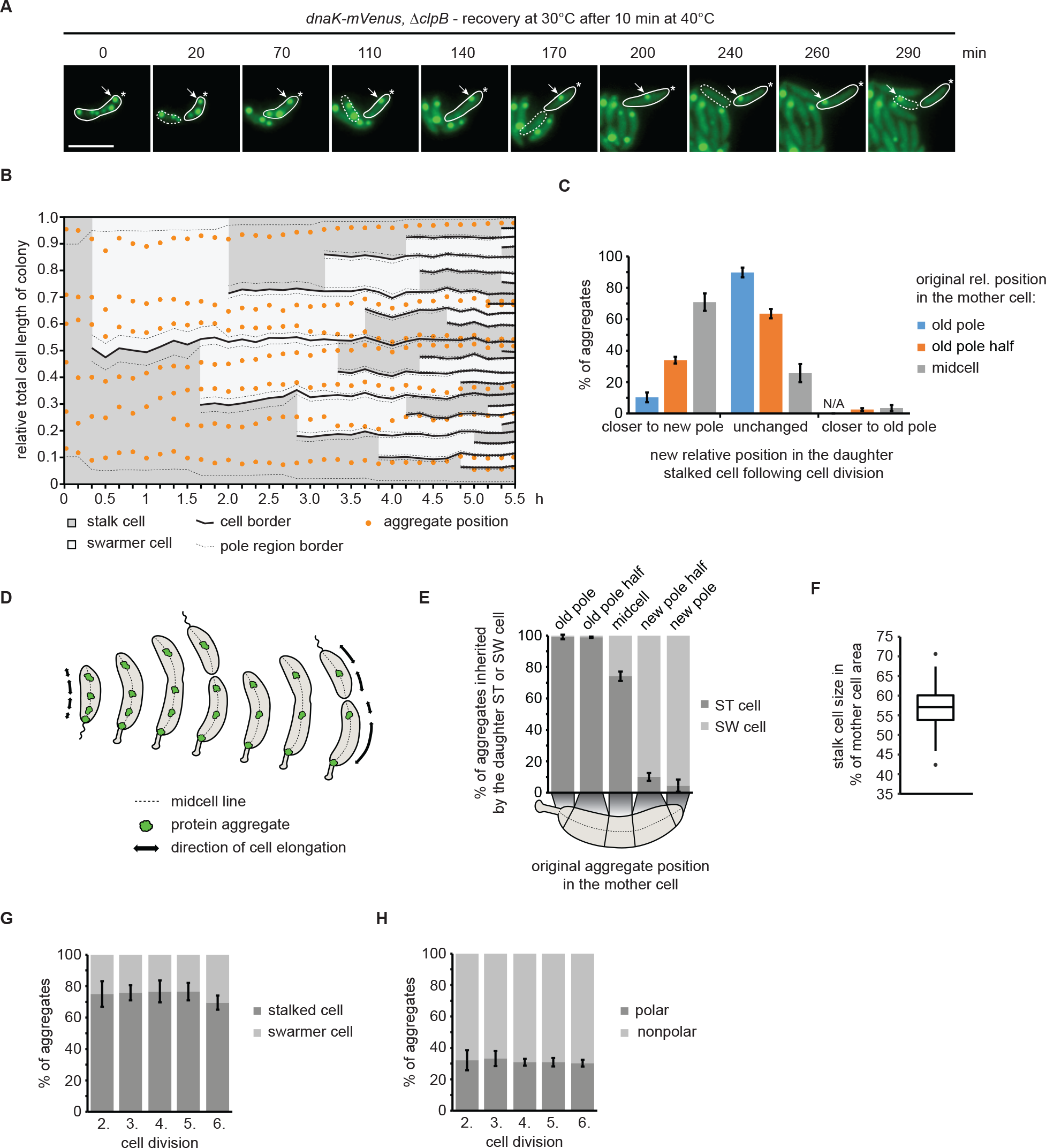
Inheritance of persistent protein aggregates in the Δ*clpB* background recovering from heat stress. (A) Representative fluorescence time-lapse images of the recovery at 30°C after exposure to 40°C for 10 min pointing out the inheritance of an aggregate originally localized in the old pole cell half after stress exposure (arrow). The star represents the location of the old pole. The stalked cell is outlined by a solid and the swarmer cell by a dashed line. (B) Kymograph normalized to the summed length of all descendants, showing relative aggregate localization and cell polarity changes during microcolony growth. Cell borders arising through cell divisions are represented by solid lines while the 10% of the cell length defined as the pole region is indicated by dashed lines. (C) Quantitative data showing how aggregates change their relative cellular position in the daughter stalked cell as a consequence of cell division. (D) Schematic model of how cell growth contributes to the movement of stationary aggregates. (E) Proportion of aggregates inherited by either a swarmer or a stalked cell as function of their cellular position in the mother cell. (F) Quantification of the amount of the mother cell area inherited by the daughter stalked cell after a division event. Quantification based on 160 division events occurring in the third and fourth generation of 33 microcolonies. (G) Percentage of aggregates inherited by the stalked or the swarmer daughter cell after the second to the sixth division. (H) Localization of aggregates tracked from the second to the sixth division. Aggregate position quantifications in (C, E, G and H) resulted from tracking the same population of aggregates from the second to the sixth division. Quantifications are based on biological triplicates for which at least 94 aggregates in at least 20 microcolonies each were tracked. For the quantifications in (C) and (E) the aggregate positional changes after each division were binned leading to at least 376 aggregate positional changes tracked per replicate. Error bars represent standard deviations.

The result that cell growth and the establishment of new cell boundaries during division frequently resulted in aggregates becoming closer to the new pole suggested that with consecutive divisions, these aggregates would eventually be inherited by a swarmer cell. Consistent with this hypothesis, we found that the cellular position in the mother cell largely determines to which daughter cell aggregates are distributed (Fig. 6E). Nearly all aggregates at the new pole or in the new pole half were inherited by the daughter swarmer cell, while essentially all aggregates at the stalked pole or in the old pole half were inherited by the daughter stalked cell (Fig. 6E). Aggregates with an original location around midcell were either inherited by the daughter stalked (74%) or swarmer (26%) cells. The preference towards the stalked cell may be attributable to the size difference between stalked and swarmer cells (Fig. 6F).

In sum, our data show that the cellular positioning of aggregates in *C. crescentus* is largely governed by the elongation of the cell and the placement of the division plane, and that the cellular position of aggregates in a mother cell determines the likelihood of being inherited by a particular cell type. As such, it is unlikely that *C. crescentus* accumulates aggregates in the old pole-inheriting daughter cell, as previously shown for *E. coli* and *Mycobacterium* (Lindner et al., 2008; Vaubourgeix et al., 2015; Winkler et al., 2010). To test this idea more directly, we determined the percentage of total aggregates that are distributed to either the old pole-inheriting stalked or the new pole-inheriting swarmer cell through five division events. Consistently, we found that the fraction of aggregates inherited by either cell type was stable (Fig. 6G). Likewise, the fraction of polar aggregates remained stable at approximately 30% over multiple generations (Fig. 6H). Stalked cells always inherited more aggregates than swarmer cells (70%) (Fig. 6G), which we attribute to the size difference and the observation that it is the stalked cell that inherits the old pole (Fig. 6C, F).

Collectively, our data show that persistent protein aggregates in *C. crescentus* are neither collected at the old pole nor asymmetrically distributed to only the stalked cell; rather, both daughter cells inherit aggregates at a constant ratio. In addition to our analysis of the Δ*clpB* background, we also investigated how persistent aggregates (Fig. 4D) are inherited in wild type cells during sustained stress at 40°C and during recovery from 44°C (Supporting Information Fig. 6, 7). These experiments yielded similar results to that obtained in the Δ*clpB* strain (Fig. 6), confirming that under these conditions *C. crescentus* does not rely on asymmetric inheritance of protein aggregates towards one cell type.

## Discussion

Our study describes the dynamics of protein aggregate formation and clearance in response to antibiotic and heat stress in the asymmetrically dividing bacterium *C. crescentus*. We find that while utilizing the same key players for aggregate clearance as other bacteria, how aggregates form in *C. crescentus* and how they are partitioned during division differs from previously described bacteria.

### Subcellular localization of aggregate clusters

We demonstrate that multiple foci of protein aggregation form in *C. crescentus* throughout the cell volume in response to both antibiotic and heat stress. In *Mycobacterium* protein aggregates also form as multiple distributed foci, although these are collected at the cell pole within one doubling time to form a pattern similar to that of *E. coli*, where aggregates rapidly localize at the poles after formation (Coquel et al., 2013; Lindner et al., 2008). In *E. coli*, it has been demonstrated that the condensed nucleoid governs collection of bigger aggregates through macromolecular crowding, enforcing movement towards and deposition at the poles (Coquel et al., 2013; Winkler et al., 2010). Compared to *E. coli* (Winkler et al., 2010), *C. crescentus* shows a more relaxed arrangement of the chromosome, where the nucleoid fills the entire cell volume (Fig. 1G). We suspect that this pattern might allow aggregate deposition throughout the entirety of the cell as opposed to only at the poles. Other large subcellular structures, such as polyphosphate granules, have been demonstrated to occupy particular locations within the cell (Henry and Crosson, 2013), however our analysis did not reveal that aggregates were more or less associated with particular cellular positions.

As cells begin to grow, whether in the presence of constant sublethal stress or during recovery from acute heat shock, we found that the majority of persistent aggregates, while static, are displaced in proportion to the elongation of the cell. We demonstrate that this displacement is continuous, although cell division may dictate a new relative position within the daughter cell. As heat shock-induced aggregates are very large and can nearly span the entire width of the cell, we expect that these may be resistant to movement within the cell, and would be largely unaffected by the movement of smaller, more mobile cellular components. Alternatively, the incorporation of un- or misfolded membrane anchored proteins into aggregate clusters may restrict intracellular movement, as has been demonstrated in *E. coli* when aggregation-prone luciferase attached to a membrane anchor is expressed (Winkler et al., 2010). In support of this latter hypothesis, we detected several membrane-associated cellular proteins to be enriched in the aggregate fraction during severe heat stress, suggesting a potential means of “tethering” the aggregate and restricting its diffusion throughout the cell.

### Aggregate clearance in response to different stress intensities

While the pattern of aggregate formation in *C. crescentus* was similar throughout all tested stress conditions, we found that the mechanisms by which aggregates are removed from individual cells largely depend on the type and intensity of stress. When recovering from exposure to sublethal stress, aggregates are rapidly dissolved by the combined activity of DnaK and ClpB resulting in complete aggregate clearance from the population within one generation. Furthermore, we found that *C. crescentus* is able to cope with continuous exposure to 40°C by rapidly upregulating the heat shock response following temperature upshift, leading to a new homeostasis of aggregate formation and dissolution, with more persistent aggregates being diluted through continuous cell growth. Noticeably, although cells incubated at 40°C grow at a similar rate as unstressed cells, they become elongated, indicating defects or delays in completing the cell cycle. A previous study showed that the cell cycle regulator CtrA is downregulated under this condition (Heinrich et al., 2016).

Our data show that when *C. crescentus* is subjected to intense stress, the persistent aggregates that form cannot be dissolved by the major chaperones, indicating that under these conditions the capacity of the protein quality control machinery is exhausted. The only way to reduce these persistent aggregates in individual cells is to distribute them during the process of cell division. Consequently, only growing and dividing cells can remove persistent aggregates, while those cells that arrest growth maintain them. Our data demonstrate that exposure to severe stress is associated with a marked heterogeneity in the ability to return to normal growth, with only a fraction of the population returning to normal growth following release to non-stress conditions. We expect that massive aggregation under severe stress causes disruption of numerous essential growth processes, supported by previous studies which indicate that strong heat shock response induction during the severe unfolding stress of high temperatures directs cellular resources away from protein translation to survival functions (Chen et al., 2017; Santra et al., 2017; Schramm et al., 2017).

### Pattern of aggregate inheritance and its physiological consequences

Our finding that persistent aggregates do not accumulate in the stalked cell, but are instead distributed to both daughter cells at a stable ratio, unveils that the pattern of aggregate inheritance strikingly differs between *C. crescentus* and the previously studied bacteria *E. coli* and *Mycobacterium*. In the latter cases, aggregates are retained only in the old pole-inheriting daughter cell, whereas the daughter cell inheriting the new pole escapes inheritance of aggregate foci (Govers et al., 2018; Lindner et al., 2008; Vaubourgeix et al., 2015; Vedel et al., 2016; Winkler et al., 2010). The resulting heterogeneity of growing microcolonies, in which one part of the population carries protein aggregates while the other part does not, has been suggested to provide a benefit for the population, either by providing a source of rejuvenation (Lindner et al., 2008; Winkler et al., 2010) or by increasing robustness to subsequent stresses (Govers et al., 2018).

An asymmetric pattern of aggregate inheritance has been a plausible, though untested, explanation for the previous observation that *Caulobacter* stalked cells decrease reproductive output with increasing cell age (Ackermann et al., 2003). While a minority of aggregates were trapped at the poles in *Caulobacter*, additional aggregates did not accumulate at this location, with the majority of cells instead eventually passing their aggregate content on to swarmer cells. Our findings suggest that rather other reasons, for example retention of older membrane components (Bergmiller et al., 2017), may underlie the phenomenon of stalked cell senescence.

Taken together, our work has revealed for the first time a shared mode of aggregate inheritance in bacteria, highlighting that asymmetric aggregate inheritance is not the sole way of managing protein aggregation. If *C. crescentus* benefits from the observed slow distribution of persistent aggregates to both daughter cell types is unclear. However, it may be advantageous to preserve both cell types in the population rather than sacrifice one type for the other in the ever-changing aquatic habitat of this bacterium.

## Materials and Methods

### Growth conditions

*C. crescentus* strains were routinely grown at 30°C in liquid PYE medium in either an Infors HT Multitron Standard or Infors HT Ecotron rotating shaker set to 200 rpm, or on solid PYE agar. Prior to analysis, all liquid cultures were grown undisturbed in exponential phase at 30 °C for three hours to allow any aggregation from other sources to be resolved or diluted. Heat shock experiments were performed by moving 100 ml flasks of 10 ml cultures growing at 30°C to a shaking incubator pre-heated to the desired temperature. Media were supplemented with the following additives when indicated; xylose 0.3 %, vanillate 500 μM. Antibiotics were used at following concentration (concentration in liquid/solid media as μg/mL): kanamycin 5 (2.5 in the case of KJ956 and KJ957)/25, spectinomycin 25/400, gentamycin 0.625/5, tetracycline 1/2, rifampicin 2.5/5. Experiments were generally performed in the absence of antibiotic when using strains in which the resistance cassette was integrated into the chromosome. *E. coli* strains for cloning purposes were grown in LB medium at 37°C, supplemented with antibiotics at following concentrations: kanamycin 30/50, spectinomycin 50/50, gentamycin 15/20, tetracycline 12/12, rifampicin 25/50. Details on strain and plasmid construction are reported in Supporting Information Text 1 and Supporting Information Tab. 1.

### Microscopy

To visualize cells, a 1% agarose in PYE slab was poured using a GeneFrame (Thermo Fisher Scientific) attached to a glass slide and coverslip, which was pre-warmed to 30°C prior to all experiments. At the indicated time points, a small volume of live cultures was transferred to the pad, sealed under a coverslip and transferred to the microscope for immediate visualization. A T*i* eclipse inverted research microscope with 100x/1.45 NA objective housed in a heated chamber, pre-heated to 30°C unless otherwise specified, was used to collect images. Fluorescence images were captured using excitation and emission filter cubes for mVenus (YFP), mCerulean (CFP), Hoechst 33258 (DAPI), and mCherry (Texas Red), ensuring that fluorescence levels did not exceed the dynamic range of the sensor.

To visualize the chromosome, cultures of *Caulobacter* were fixed using a final concentration of 4% formaldehyde for 5 min, following which DNA was stained for 25 min using Hoechst 33258 at a final concentration of 2 μg/mL. Fixed cells were transferred to a 1% agarose pad and visualized under the microscope as described above for live cells.

### Custom-built STED set-up and STED imaging

The super-resolution imaging has been performed at a custom-built STED set-up. The samples were fixed and adhered to slides as described by (Hiraga et al., 1998), and labelled with an anti-GFP nanobody coupled to ATTO 594 (GFP-Booster_Atto594, ChromoTek). The dye was excited with a 561 nm pulsed diode laser (PDL561, Abberior Instruments) and subsequently depleted with a 775 nm pulsed laser (KATANA 08 HP, OneFive), both operating at 40 MHz. The depletion beam was shaped to a donut in the focal plane by the use of a spatial light modulator (LCOS-SLM X10468-02, Hamamatsu Photonics). The excitation laser and depletion laser were coupled together and scanned over the sample using fast galvanometer mirrors (galvanometer mirrors 6215H + servo driver 71215HHJ 671, Cambridge Technology). The laser beams were focused onto the sample using a HC PL APO 100x/1.40 Oil STED White objective lens (15506378, Leica Microsystems), through which also the fluorescence signal was collected. After de-scanning and de-coupling of the fluorescence signal, it was put through a confocal pinhole (1.28 Airy disk units) and detected through a bandpass filter (ET615/30m, Chroma Technology) and a notch filter (ZET785NF, Chroma Technology) with a fiber-coupled APD (SPCM-AQRH-14-FC, PerkinElmer).

The imaging was done with a 561 nm excitation laser power of 1.1-8.4 μW and a 775 nm depletion laser power of 132-158 mW, both measured at the first conjugate back focal plane of the objective. The pixel size for all images was set to 25 nm. Each image was acquired by adding up 10 line scans for each line, each with a pixel dwell time of 20 μs, resulting in a total pixel dwell time of 200 μs.

### Aggregate size analysis

The aggregate sizes were calculated from groups of fitted line profiles in the STED and confocal images. Using Fiji (ImageJ), aggregates were picked out and line profiles were drawn across them. The line profiles were extracted and fitted with a Lorentzian function, from which the full width at half maximum was extracted as the aggregate size. The number of aggregates used for each mean and standard deviation calculation was at least 60.

### Image and statistical analysis

To prepare the appropriate image formats and to perform basic analyses such as foci enumeration and generation of line profiles, the Fiji software package was used. Information on the area of individual cells, their fluorescence intensity, as well as their corresponding aggregate number and position in Figure 1D, E, 4, 5 and Supplemental Information Fig. 2 and 4 were gathered by analyzing binary masks of cells and aggregate foci using the ImageJ package MicrobeJ (Ducret et al., 2016). The Oufti software package (Paintdakhi et al., 2016) was used to generate data on population cell lengths in Figure 4F. To determine fluorescence intensity per bacterial pixel shown in Figure 1I and 3C, the MATLAB package SuperSegger (Stylianidou et al., 2016) was used to perform segmentation on images followed by calculation of total cell area per image and bacterial fluorescence intensity per image. For the preparation of cell masks, bacterial segmentation was performed on phase contrast images using SuperSegger and all segmentation was manually checked before further analysis. Masks of the aggregate signal were manually prepared by labeling corresponding areas. The colony kymograph in Figure 6B was prepared based on cell and aggregate masks using SuperSegger to generate cell linking, cell length, and aggregate position measurements. Aggregate inheritance and positional changes after sequential divisions (Fig. 6, Supplemental Information Fig. 5, 6, 7) as well as the aggregate life span (Fig. 4D, G, 5F, Supplemental Information Fig. 4B) were manually tracked. Data sets generated through all image analysis programs were analyzed and visualized using MATLAB and R software packages.

### Immunoblotting

Cell pellets were collected following the indicated treatments and time points and normalized by units OD_600_ through dilution in Laemmli buffer. Diluted samples were boiled at 98°C for 10 min and loaded into Stain Free Tris-glycine gels (Bio-Rad) for separation by SDS-PAGE and transfer to a nitrocellulose membrane as per manufacturer guidelines. Membranes were blocked for 1 h in 5% skim milk powder in TBST and protein levels were detected using either anti-DnaK antibody (Schramm et al., 2017) or the anti-GFP antibody (Thermo Fisher Scientific, #A-11122). A secondary anti-rabbit HRP-conjugated antibody was used to detect primary antibody binding, and SuperSignal Femto West (Thermo Fisher Scientific) used to develop the membrane. Blots were scanned using a Chemidoc (Bio-Rad) system and signal quantification was performed using the Image Lab software package (Bio-Rad).

### Aggregation assay

The aggregation assay was performed as described in (Schramm et al., 2017). About 40 OD_600_ units of cells with an OD_600_ between 0.2-0.4 either grown at 30°C or heat stressed were harvested. For the heat treatment an exponential overnight culture grown at 30°C was pelleted (6000 g, 10 min, RT), resuspended in a flask containing 45°C pre-warmed medium and then incubated shaking at the same temperature for 1 h. A portion of the aggregate fraction was subjected to SDS-PAGE and the separated proteins were visualized by Coomassie staining (Instant Blue Protein Stain, Expedeon).

### Proteomic analysis of protein aggregates

For the identification of endogenous aggregating and aggregate associated proteins in *Caulobacter crescentus* whole protein aggregate fractions were subjected to mass spectrometry analysis. 0.1 μg horse cytochrome C were added to each sample to serve as internal protein abundance standard. The mass spectrometry analysis of the samples was performed using an Orbitrap Velos Pro mass spectrometer (Thermo Scientific). An Ultimate nanoRSLC-HPLC system (Dionex), equipped with a custom 20 cm x 75 μm C18 RP column filled with 1.7 μm beads was connected online to the mass spectrometer through a Proxeon nanospray source. 1-15 μL of the tryptic digest (depending on sample concentration) were injected onto a C18 pre-concentration column. Automated trapping and desalting of the sample was performed at a flowrate of 6 μL/min using water/0.05% formic acid as solvent. Separation of the tryptic peptides was achieved with the following gradient of water/0.05% formic acid (solvent A) and 80% acetonitrile/0.045% formic acid (solvent B) at a flow rate of 300 nL/min: holding 4% B for five minutes, followed by a linear gradient to 45% B within 30 minutes and linear increase to 95% solvent B in additional 5 minutes. The column was connected to a stainless steel nanoemitter (Proxeon, Denmark) and the eluent was sprayed directly towards the heated capillary of the mass spectrometer using a potential of 2300 V. A survey scan with a resolution of 60000 within the Orbitrap mass analyzer was combined with at least three data-dependent MS/MS scans with dynamic exclusion for 30 s either using CID with the linear ion-trap or using HCD combined with orbitrap detection at a resolution of 7500. Data analysis was performed using Proteome Discoverer (Thermo Scientific) with SEQUEST search engine using Uniprot databases. The peptide peak areas of each protein in a sample were normalized to that of the internal Cytochrome C standard. Proteins were considered to be specifically enriched in the aggregate fraction if they were only detected in the heat-treated sample or more abundant than in the control.

### Spot colony formation assays

Cultures with an OD_600_ between 0.1-0.4 were diluted to 0.05 after which a 1:10 serial dilution was performed. 2 μL of each dilution were spotted on PYE agar plates.

## Acknowledgements

We thank Dr. Claes Andréasson and members of the Jonas lab for helpful discussion. We also thank Dr. Uwe Linne and the mass spectrometry facility at the University of Marburg for their assistance. The study was financially supported by funding from the LOEWE program of the state Hessen, funding of Strategic Research Areas (SFO) from Stockholm University, and a future research leaders grant from the Swedish Foundation for Strategic Research (SSF).

## Supporting Information

### Content

**Supporting Information Figure 1.**
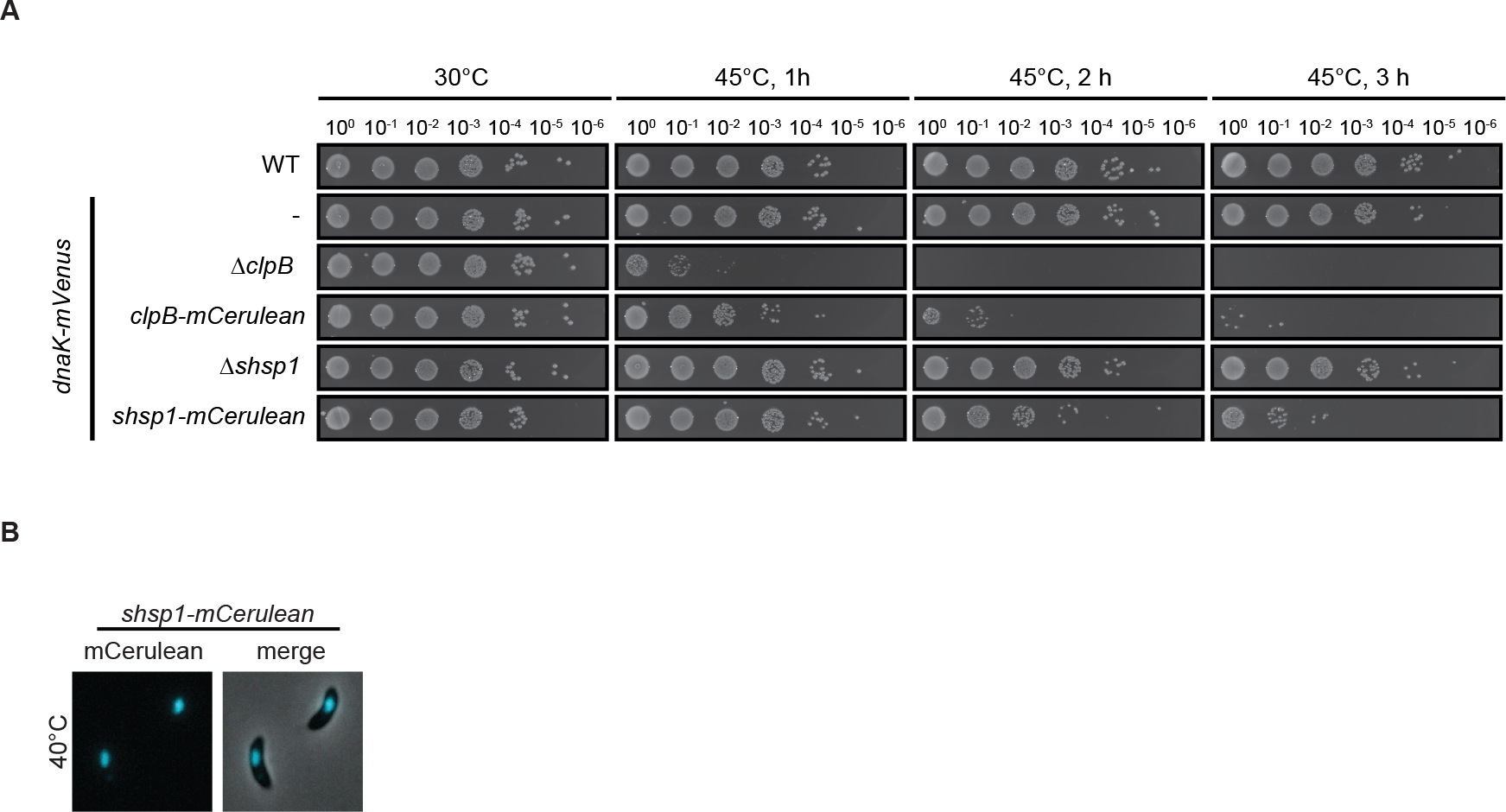
Viability and functionality of the fluorescent reporters used in this study. (A) Spot assays showing resistance to exposure of 45°C of the wild type and strains bearing the indicated deletions or fusions in addition to DnaK-mVenus. (B) Microscopy images demonstrating localization of the small heat shock proteins sHSP1 after 1 h at 40°C.

**Supporting Information Figure 2.**
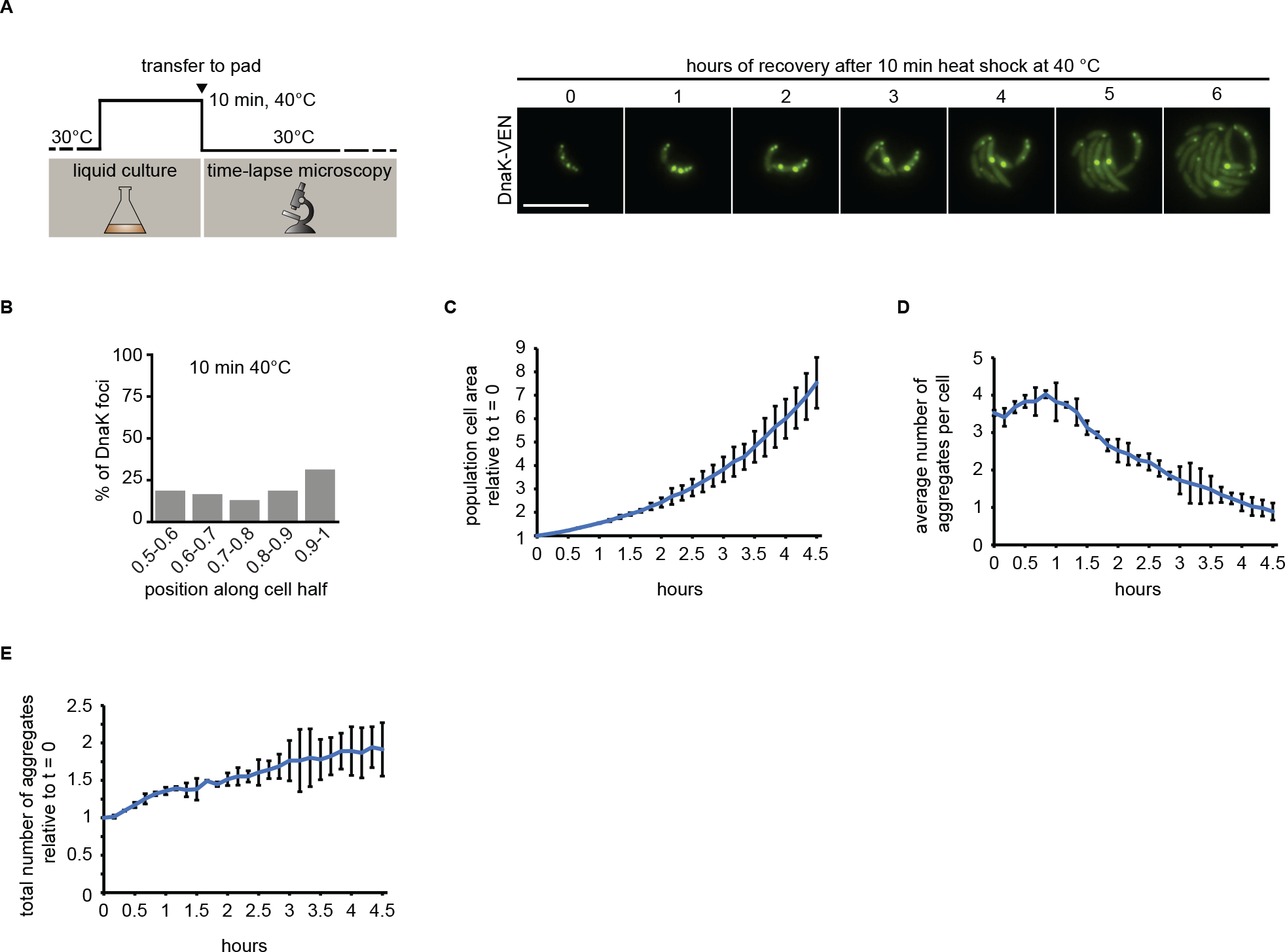
Δ*clpB* cells recovering from mild heat stress do not dissolve aggregates. (A) Schematic of recovery experiment (left) and representative images from Δ*clpB* cell recovery after exposure to 40°C for 10 min. (B) Graph illustrating the position of DnaK-mVenus foci along five bins from midcell (0.5) to the cell pole (1) in Δ*clpB* cells after a 10 min heat shock at 40°C in liquid medium. Quantification based on 313 cells harboring 1056 aggregates. (C) Total population cell area increase relative to the area after stress exposure (t=0). (D) Average number of aggregates per cell over time. (E) Total number of aggregates present in the population over time normalized to time=0. Quantifications in (B-D) show the means of biological duplicates for which 19 microcolonies each were quantified. Error bars represent the standard deviation.

**Supporting Information Figure 3.**
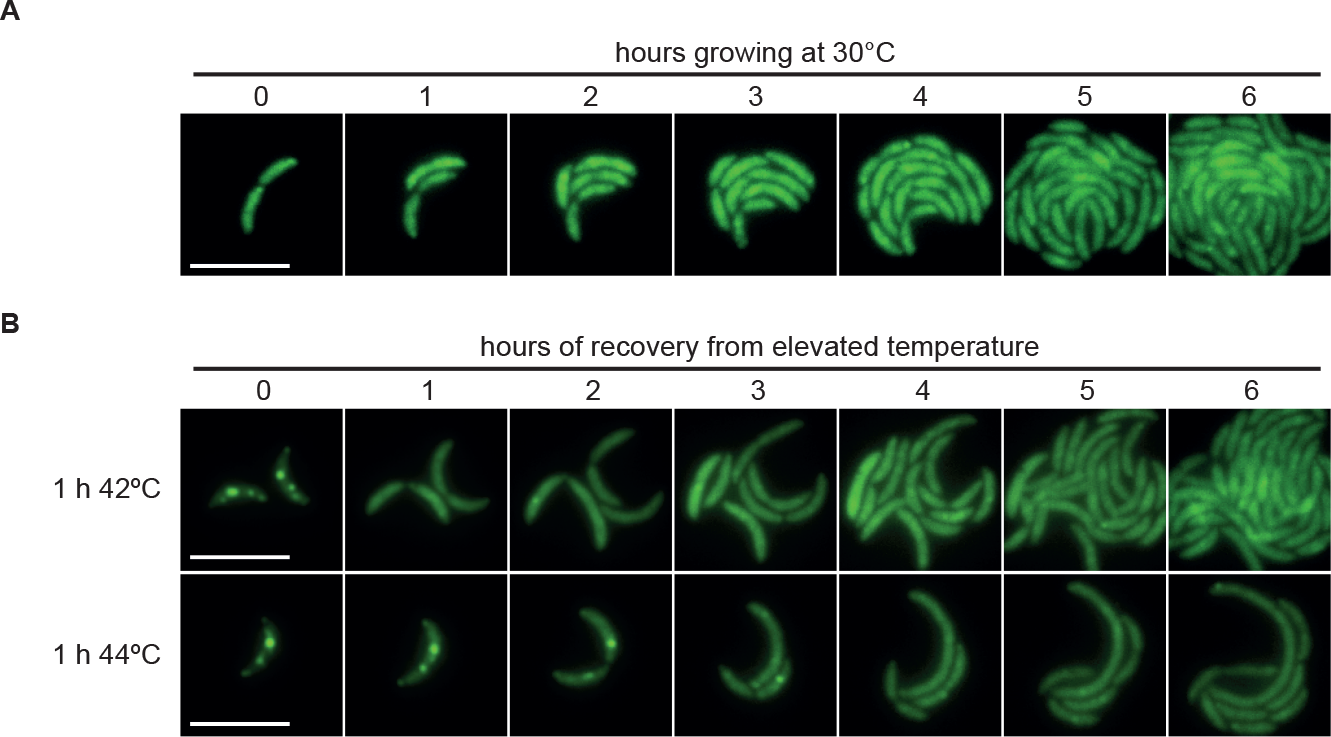
Representative time courses of cells continuously growing at 30°C or recovering from elevated temperature. (A) Representative images of *C. crescentus* grown at 30°C in a heated microscopy chamber over a six hour period. (B) Representative images of *C. crescentus* grown at 30°C in a heated microscopy chamber during recovery from exposure to 42°C or 44°C over a six hour period. Scale bar is 5 μm.

**Supporting Information Figure 4.**
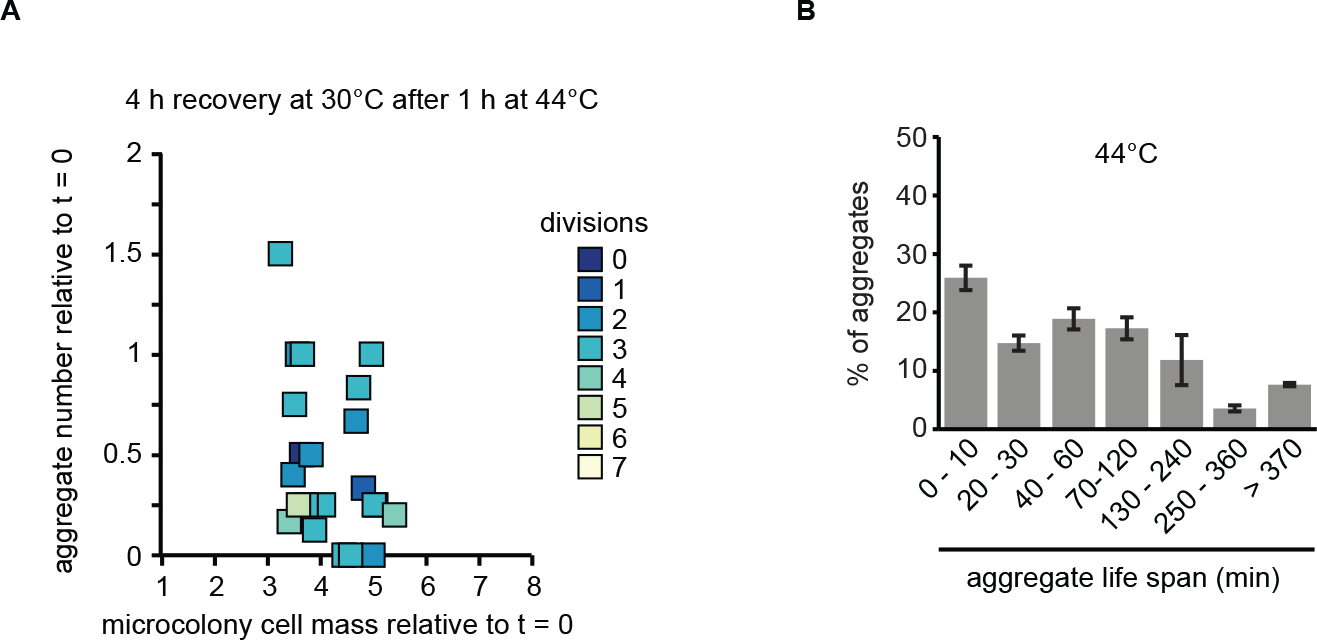
Growth and aggregate clearance in cells recovering from 1 h of exposure to 44°C. (A) Scatter plot representing relative area increase, relative aggregate number normalized to the amount present at stress release, and the number of divisions of 22 cells and their progeny after four hours (the time at which the average area of the population has quadrupled) at 30°C after release from one hour of exposure to 44°C. (B) Quantification of aggregate life span in cells recovering from 44°C. Aggregates present or emerging in the first 400 min of a fluorescence time-lapse movie were tracked until 500 min. Images were acquired every 10 min. Quantifications show the means of biological duplicates for which at least ten microcolonies and 94 aggregates each were analyzed. Error bars represent standard deviations.

**Supporting Information Figure 5.**
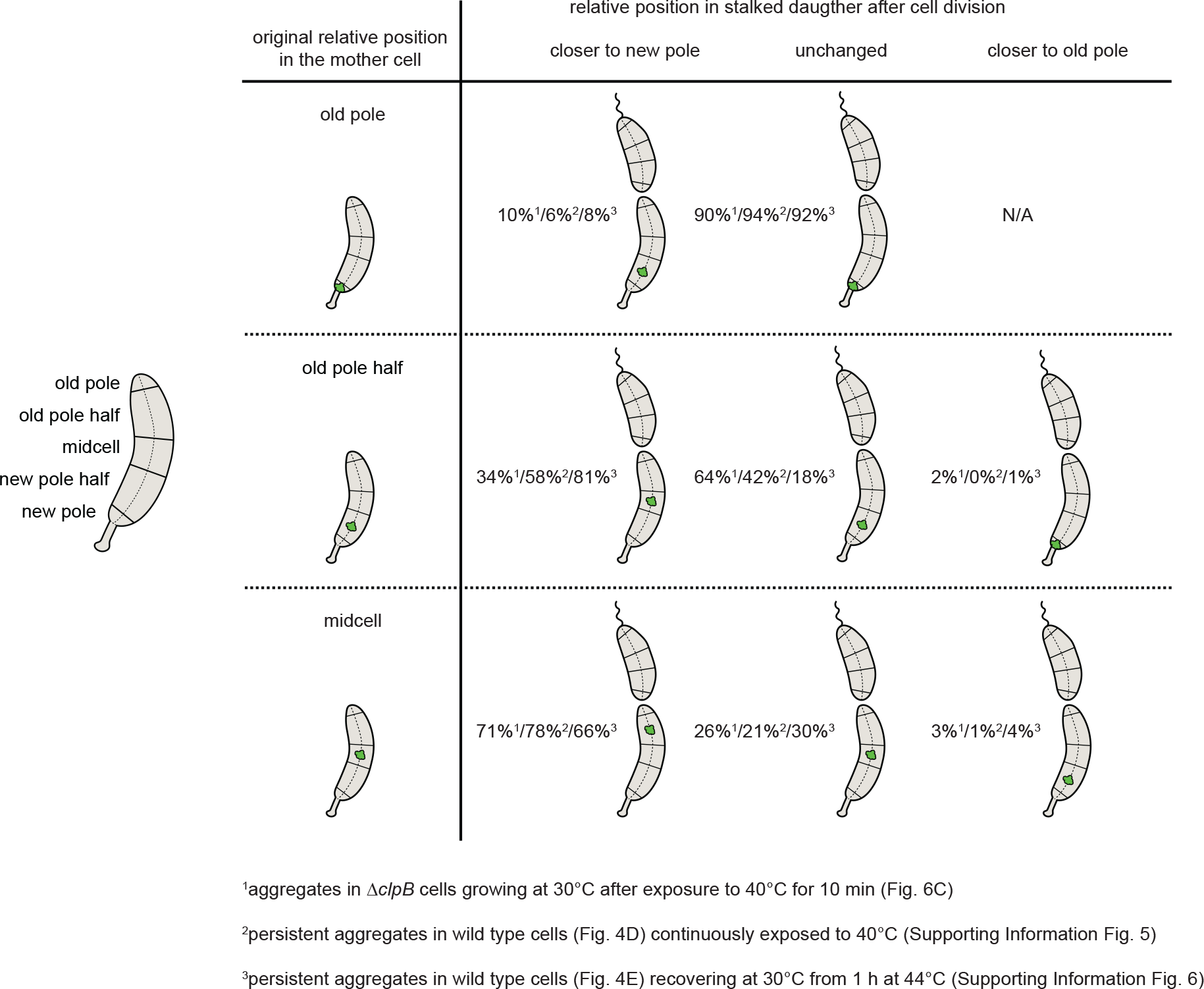
Schematic table summarizing how positional changes after cell divisions ranked by the position in the mother cell were quantified. The table shows how aggregate inheritance was analyzed for generating Figure 6C and Supporting Information Fig. 6A, 7A. Percentage numbers represent the distribution of aggregate positional changes.

**Supporting Information Figure 6.**
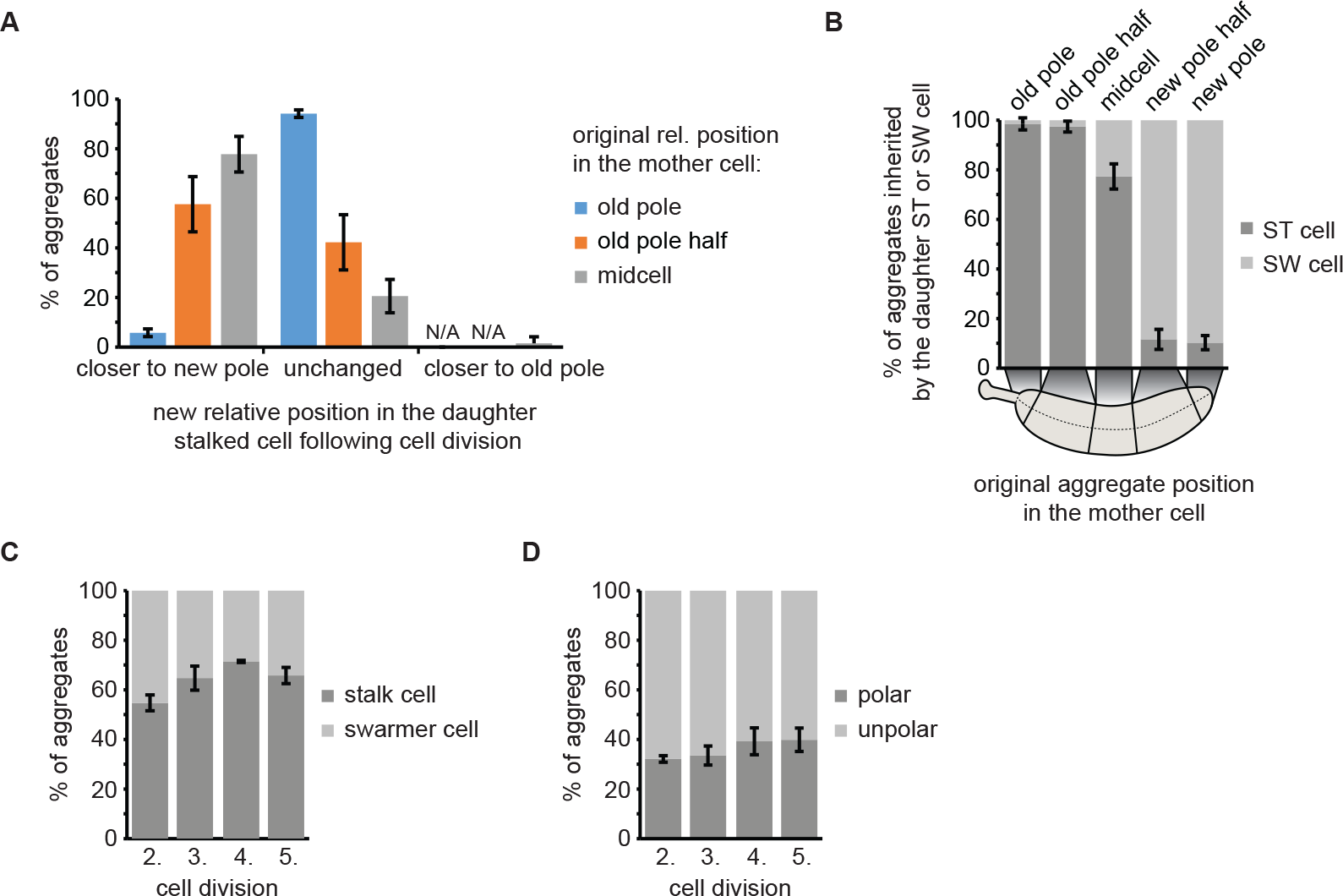
Inheritance of persistent protein aggregates in DnaK-mVenus cells continuously exposed to 40°C. (A) Distribution of aggregates becoming closer to the new pole, remaining stationary or becoming closer to the old pole in the daughter stalked cell, ranked by position in the mother cell. (B) Proportion of aggregates inherited by either a swarmer or a stalked cell as function of their cellular position in the mother cell. (C) Percentage of aggregates inherited by the stalked or the swarmer daughter cell after the second to the fifth division. (D) Localization of aggregates tracked from the second to the fifth division. Aggregate position quantifications resulted from tracking the same population of aggregates from the second to the fifth division. Quantifications are based on biological triplicates for which at least 82 aggregates in at least 21 microcolonies each were tracked. For the quantifications in (A) and (B) the aggregate positional changes after each division were binned leading to at least 246 aggregate positional changes tracked per replicate. Error bars represent standard deviations.

**Supporting Information Figure 7.**
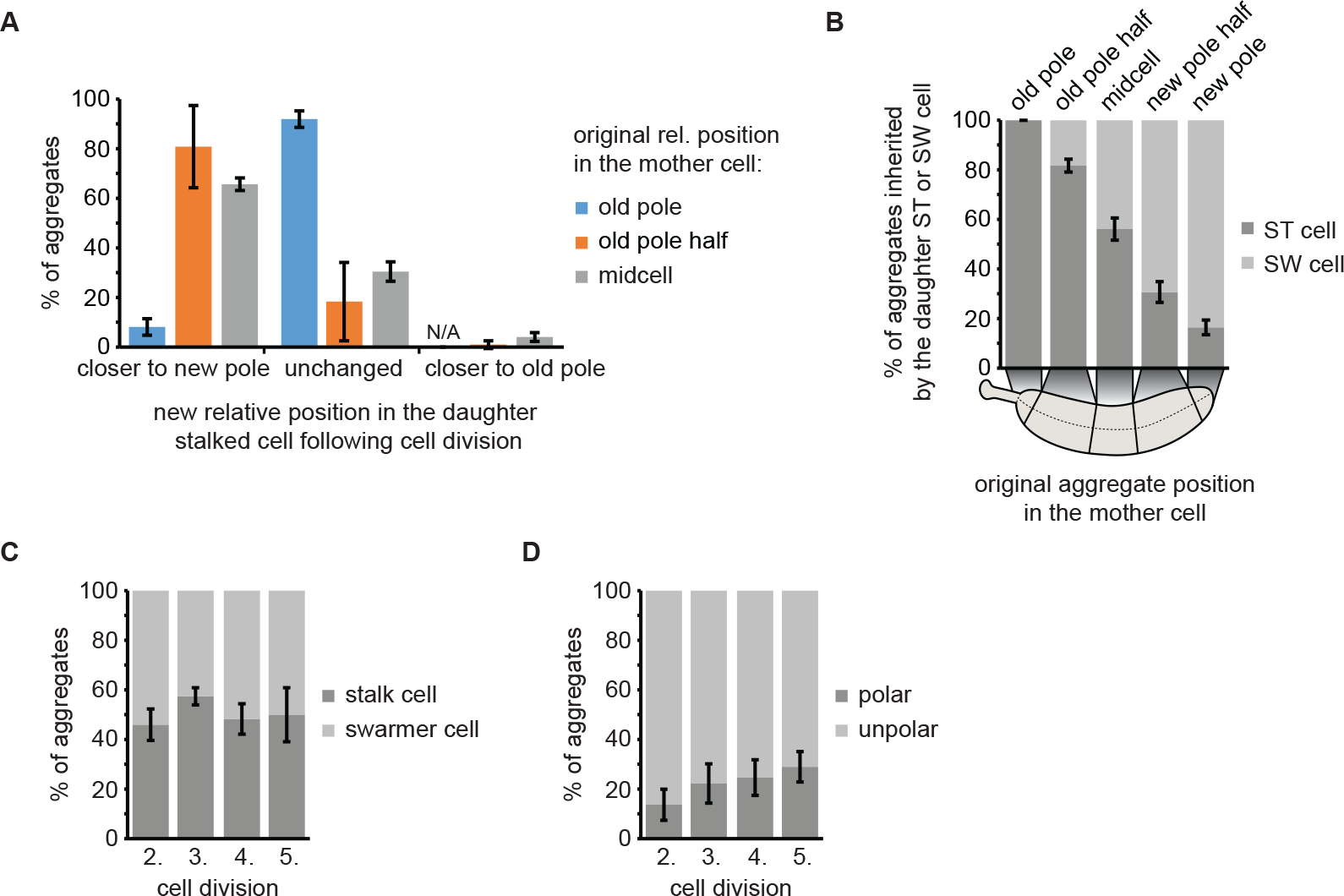
Inheritance of persistent protein aggregates in DnaK-mVenus cells continuously exposed to 44°C. (A) Distribution of aggregates becoming closer to the new pole, remaining stationary or becoming closer to the old pole in the daughter stalked cell, ranked by position in the mother cell. (B) Proportion of aggregates inherited by either a swarmer or a stalked cell as function of their cellular position in the mother cell. (C) Percentage of aggregates inherited by the stalked or the swarmer daughter cell after the second to the fifth division. (D) Localization of aggregates tracked from the second to the fifth division. Aggregate position quantifications resulted from tracking the same population of aggregates from the second to the fifth division. Quantifications are based on biological triplicates for which at least 96 aggregates in at least 85 microcolonies each were tracked. For the quantifications in (A) and (B) the aggregate positional changes after each division were binned leading to at least 288 aggregate positional changes tracked per replicate. Error bars represent standard deviations.

### Additional Supporting Information

**Supporting Information Table 1.** Strains, plasmids and primers used in this study

**Supporting Information Table 2.** Proteins enriched in the aggregated fraction in *C. crescentus* following exposure to 45°C

**Supporting Information Movie 1.** Time lapse microscopy of DnaK-mVenus upshift to 40°C

## Supporting Information Text 1

### Extended description of plasmid construction

#### Construct for fluorescence reporter tagging of *dnaK* at the native locus (pKJ936)

A fragment containing the upstream homology region (UHR) made up of the last 600 bp of *dnaK* before the stop codon and encoding a GSG linker at the 3’-end was amplified using OFS193/318 from chromosomal DNA. The downstream homology region (DHR) encompassing the 665 bp downstream of the gene was amplified with OFS197/198 from chromosomal DNA. The fluorescence reporter encoding gene *mVenus* was amplified with OFS289/307. The fragments were assembled into *Hind*III/*Eco*RI restricted pNPTS138 by Gibson assembly (Gibson et al., 2009).

#### Construct for fluorescence reporter tagging of *clpB* at the native locus (pKJ937)

A fragment comprising the UHR encompassing the last 635 bp of *clpB* before the stop codon as well as encoding a GSG linker at the 3’-end was amplified using OFS287/288 from chromosomal DNA. The DHR containing the 601 bp downstream of *clpB* was amplified with OFS290/291. The fluorescence reporter encoding gene *mCerulean* was amplified with OFS289/307. The fragments were assembled into OFS285/286 amplified pNPTS138 by Gibson assembly.

#### Construct for fluorescence reporter tagging of *shsp1* (*CCNA_02341*) at the native locus (pKJ938)

A fragment comprising the UHR encompassing 223 bp upstream of *shsp1* (CCNA_02341), the entire gene except for the stop codon and encoding a GSG linker at the 3’-end was amplified using OFS292/293 from chromosomal DNA. The DHR containing the 594 bp downstream of *shsp1* was amplified with OFS294/295. The fluorescence reporter encoding gene *mCerulean* was amplified with OFS289/307. The fragments were assembled into OFS285/286 amplified pNPTS138 by Gibson assembly.

#### Constructs for vanillate-dependent expression of mCherry-ELK16 from the chromosomal *vanA* locus (pKJ939)

For the construction of pKJ939 the TP-linker-ELK16 encoding sequence was added to the 3’-end of *mCherry* by sequential PCRs in two steps. First *mCherry* was amplified using OFS308/309 and the resulting fragment was used as a template for an amplification with OFS308/310. The resulting fragment was restriction cloned into *Nde*I/*Xba*I cut pBVMCS-2 resulting in pKJ949. This construct was used as a template to amplify *mCherry-ELK16* which was then restriction cloned into *Nde*I/*Sac*I cut pVCHYN-4 resulting in pKJ939.

#### Constructs for xylose-dependent expression of mCerulean-tagged endogenous aggregating proteins from the chromosomal *xylX* locus (pKJ941-943)

For the construction of pKJ941, pKJ942 and pKJ943, the endogenous genes were amplified from chromosomal DNA using OFS865/866, OFS869/870 or OFS880/881, respectively, and assembled with either OFS867/868, OFS871/868 or OFS875/879 amplified *mCerulean* into *Nde*I/*Sac*I cut pXCHYN-1 by Gibson assembly.

#### Constructs for deleting chaperone and protease encoding genes (pKJ944-946)

For making pKJ944 the UHR containing the 608 bp upstream and the first 15 bp of *ibpA* was amplified from genomic DNA using OFS795/796. The DHR containing the last 27 bp and the 586 bp downstream of *ibpA* was amplified with OFS797/798. The fragments were assembled with an OFS801/802 amplified *rif^R^*-cassette into *Eco*RI/*Hind*III cut pNPTS138 by Gibson assembly. In order to construct pKJ945 the UHR encompassing the 646 bp upstream and the first 15 bp of *ibpB* was amplified from genomic DNA using OFS809/817. The DHR containing the last 27 bp and the 612 bp downstream of *ibpB* was amplified using OFS811/818. The homology region were assembled with an OFS25/26 *tet^R^*-cassette (pNPTS-*lon::tet^R^*) (Leslie et al., 2015) into *Eco*RI/*Hind*III cut pNPTS138 by Gibson assembly. For the generation of pKJ946 the UHR comprising the 606 bp upstream and the first 15 bp of *clpB* was amplified from genomic DNA using OFS803/807. The DHR encompassing the last 27 bp and the 537 bp downstream of *clpB* was amplified with OFS805/808. The homology region were assembled with an OFS25/26 amplified *tet^R^*-cassette (pNPTS-*lon::tet^R^*) (Leslie et al., 2015) into *Eco*RI/*Hind*III cut pNPTS138 by Gibson assembly.

#### Constructs for ectopic overexpression of *dnaK*- and *dnaK(K70A)-GSG-mVenus* (pKJ950-951)

For constructing pKJ950 *dnaK* was amplified from chromosomal DNA with OFS302/303 and assembled with OFS289/307 amplified *mVenus* into the OFS300/320 amplified vector pBVMCS-2 by Gibson assembly. In case of pKJ951 the mutation in *dnaK* was introduced by assembling two fragments by Gibson assembly which harbor the desired sequence alteration in the overlapping region. The fragments were amplified using OFS306/320 and OFS305/307, respectively, using pKJ951as template.

### Extended description of strain construction

#### Fluorescence reporter tagging of DnaK, ClpB and sHSP1 (CCNA_02341) at the native locus (KJ952, KJ953, KJ969)

C-terminally tagging chromosomal chaperone genes with fluorescent reporter encoding sequences was achieved by a two-step recombination procedure (Skerker et al., 2005). *C. crescentus* NA1000 was transformed with pKJ936 to generate KJ952 or with pKJ948 to generate KJ969. KJ952 was transformed with pKJ937 to generate KJ953. First integrants were selected for by plating on kanamycin-containing plates. Selected integrants were grown over night in PYE medium lacking kanamycin and plated on 3 % sucrose containing plates. Clones being both sucrose-resistant and kanamycin sensitive were verified for plasmid excision and fluorescent reporter gene insertion at the correct locus by colony PCR and fluorescence microscopy.

#### Vanillate- and xylose-dependent expression of fluorescently tagged artificial and endogenous aggregating proteins as well as DnaK-GSG-mVenus from the chromosome (KJ955, KJ959-961)

Plasmids encoding for fluorescently tagged artificial, endogenous aggregating and untagged fluorescent proteins under the control of *P_vanA_* or *P_xylX_* were integrated through homologous recombination at the chromosomal *vanA* or *xylX* site, respectively. Integrants were selected by the plasmid encoded antibiotic resistance and verified by colony PCR. KJ953 was transformed with pKJ939 to generate KJ955. For the generation of KJ959, KJ960 and KJ961, KJ952 cells were transformed with pKJ941, pKJ942 and pKJ943, respectively.

#### Wt DnaK- and DnaK(K70A)-GSG-mVenus overexpressing strains (KJ956, KJ957)

*C. crescentus* NA1000 was transformed with the replicating plasmids pKJ950 and pKJ951 to obtain KJ956 and KJ957, respectively.

#### Chaperone and protease knockout strains (KJ962-966)

Antibiotic-resistance marked knockouts of chaperone and protease encoding genes were obtained by two step recombination under constant exposure to the antibiotic against which the resistance cassette replacing the deleted gene sequence provides protection. KJ952 was transformed with either pKJ944, pKJ945, pKJ946 or pNPTS-*lon::tet^R^* for the generation of KJ962, KJ963, KJ964 and KJ966, respectively. For constructing KJ965, KJ962 was transformed with pKJ945. Clones were verified by colony PCR.

